# Human ASPDH is a 2-aminomuconate reductase that produces L-2-aminohex-3-enedioic acid in tryptophan catabolism

**DOI:** 10.64898/2026.06.05.730449

**Authors:** Giulia Sassi, Giulia Mori, Serena Facchetti, José Fernando Rinaldi de Alvarenga, Marina Simona Robescu, Elena Dembech, Marco Vezzoli, Giovanni Merici, Marco Malatesta, Sara Dobani, Daniele Del Rio, Letizia Bresciani, Giacomo Quilici, Davide Cavazzini, Roberto Battistutta, Francesco Sansone, Andrea Armirotti, Roberto Ferrari, Pedro Mena, Daniela Ubiali, Alessio Peracchi, Claudio Rivetti, Riccardo Percudani

**Affiliations:** Department of Chemistry, Life Sciences and Environmental Sustainability, Università di Parma, 43124 Parma, Italy; Human Nutrition Unit, Department of Food and Drug, Università di Parma, 43124 Parma, Italy; Department of Drug Sciences, Università degli Studi di Pavia, 27100 Pavia. Italy; Centro di Servizi e Misure “Giuseppe Casnati”, Università di Parma, 43124 Parma, Italy; Department of Chemical Sciences, Università degli Studi di Padova, 35131 Padova, Italy; Fondazione Istituto Italiano di Tecnologia, 16163 Genova, Italy

## Abstract

Most tryptophan catabolism in animals occurs through the kynurenine pathway, which generates the essential NAD cofactor and multiple bioactive metabolites. Knowledge of this pathway in eukaryotes ends at the unstable intermediate 2-aminomuconate (2-AM). Here, by leveraging evolutionary information from more than 5,000 eukaryotes, we identify two distinct genes acting downstream of 2-AM in fungi and metazoa. The fungal gene is homologous to bacterial 2-AM deaminase, whereas the metazoan gene is homologous to aspartate dehydrogenase (ASPDH), which in prokaryotes catalyses the first reaction of NAD biosynthesis. Biochemical and structural analyses show that human ASPDH has evolved an unprecedented function as an NAD(P)H-dependent 2-AM reductase (AMR) in tryptophan catabolism. The reaction forms L-2-aminohex-3-enedioic acid, an unsaturated α-amino acid absent from current biological databases. Isotope-labeling NMR experiments and structural modelling support a mechanism in which hydride transfer is coupled to double-bond rearrangement of the conjugated system. These findings reveal a previously unknown metazoan branch of the kynurenine pathway, expand the repertoire of endogenous amino acids, and illustrate how comparative genomics can uncover hidden reactions in human metabolism.

## Introduction

Among the catabolic pathways for essential amino acids, L-tryptophan (Trp) degradation has a particular relevance in animals ^1–7^. In the gut and central nervous system, Trp is the source of neurotransmitters serotonin, melatonin and other indoleamines via the serotonergic pathway ^4,8^. However, most Trp flux in humans and other mammals occurs through the kynurenine pathway (KP), particularly in the liver. This pathway provides a *de novo* route to NAD biosynthesis and generates a range of bioactive metabolites, collectively termed kynurenines^3,5^. Among these, L-kynurenine (Kyn) and kynurenic acid are immunoregulators ^9,10^, 3-hydroxykynurenine (3-HK) is a redox-active intermediate ^11^, whereas xanthurenic acid and quinolinic acid affect brain functions ^12,13^, and picolinic acid exerts anabolic effects on bone ^14^. Through these activities, kynurenines influence diseases ranging from neurological conditions to inflammation and cancer progression ^1,5,6^.

The canonical pathway (**Fig. 1a**) begins with oxidation of Trp to *N’*-formylkynurenine by tryptophan 2,3-dioxygenase (TDO2) in the mammalian liver or by indoleamine 2,3-dioxygenases (IDO1/2) in extrahepatic tissues. Subsequent deformylation by arylformamidase (AFMID) produces the metabolite that gives the pathway its name. Hydroxylation by kynurenine 3-monooxygenase (KMO) yields 3-HK, which is cleaved by kynureninase (KYNU) to alanine and 3-hydroxyanthranilate (3-HAA). This upstream segment of the pathway is diversified across tissues and lineages. In mammals, Kyn can be converted to kynurenic acid via transamination, whereas 3-HK can be converted to xanthurenic acid. In insects, cephalopods and other protostomes, 3-HK is instead the precursor of ommochromes, pigments responsible for integument and eye coloration ^15,16^. In *Caenorhabditis elegans* and other anoxia-tolerant protostomes, 3-HAA is a precursor of rhodoquinone, a lipid electron carrier that supports oxygen-independent respiration ^17,18^.

**Fig 1.**
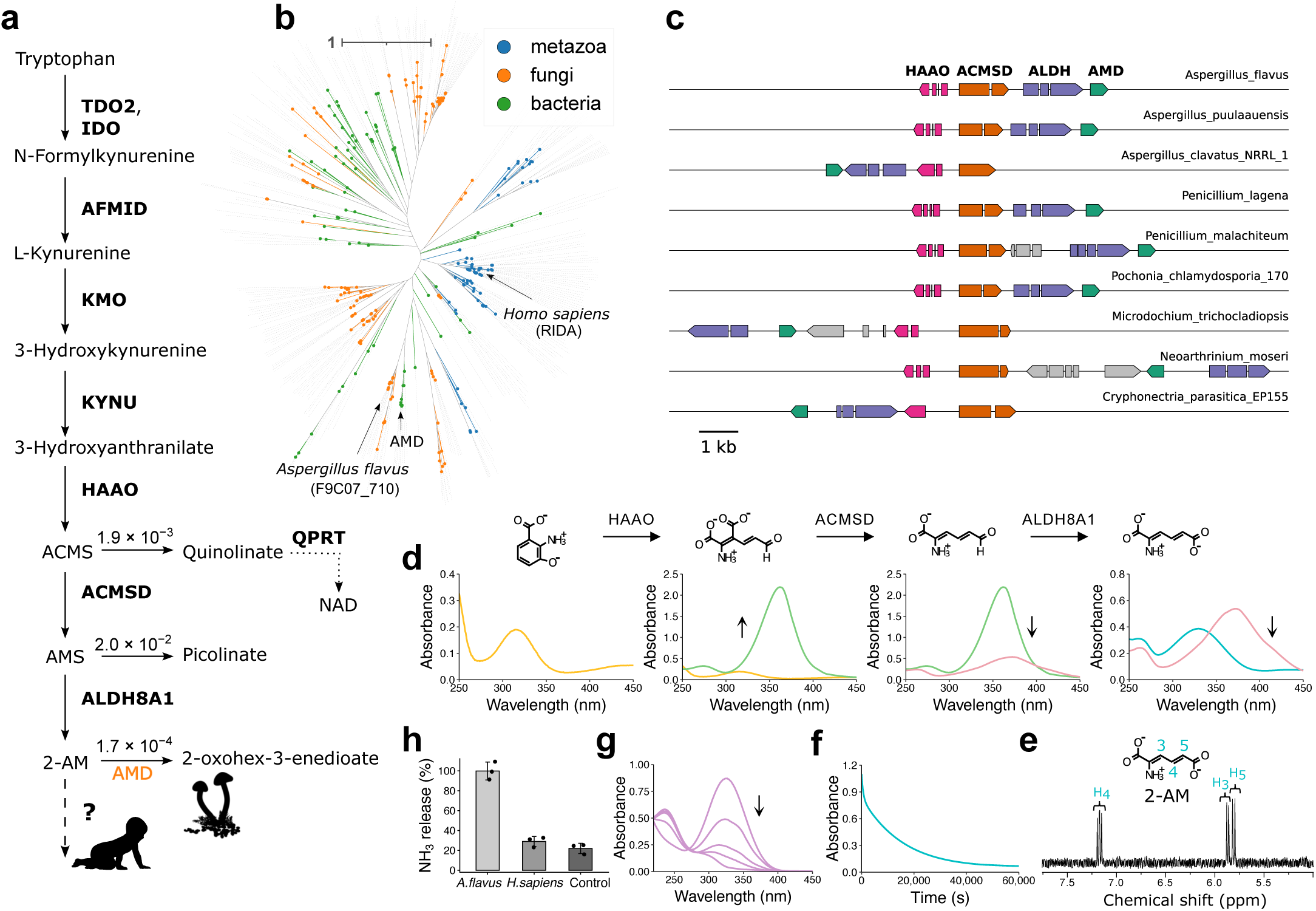
Identification of fungal AMD highlights a missing step in metazoan KP. **a.** Scheme of the KP in eukaryotes; known human genes are indicated by bold symbols; non-enzymatic rate constants (s^−1^) in near-physiological conditions are shown ^24–26^. The identified fungal AMD and the missing human gene are indicated at the last branching point. **b.** IPR006175 family tree in bacteria, fungi, and metazoa. The position of the validated bacterial AMD is indicated along with the human (RIDA) and *A. flavus* protein (F9C07_710) selected for functional characterization. **c,** Microsynteny analysis of the 2-AMD locus in different ascomycetes showing the conserved genomic association with other KP genes. **d.** Enzymatic synthesis of 2-AM, monitored by UV–visible spectroscopy; chemical structures are shown above the corresponding spectra. **e,** ^1^H NMR spectrum of purified 2-AM with proton assignment. **f**. Decay of the 2-AM absorption signal at 326 nm, consistent with spontaneous deamination. **g,** UV–visible spectra of the *A. flavus* AMD reaction with 2-AM; spectra were recorded 1 s, 15 s, 3 min, and 1 h after enzyme addition. **h,** Ammonium release from 2-AM in the presence of *A. flavus* AMD, human RIDA or no enzyme, monitored for 500 s using a GDH-coupled assay. Points show individual measurements (n = 3); bars represent the mean ± SD.

The downstream segment of the pathway links Trp degradation to NAD biosynthesis and central carbon metabolism ^19–22^. Oxidative ring opening of 3-HAA by 3-hydroxyanthranilate 3,4-dioxygenase (HAAO) produces 2-amino-3-carboxymuconate-6-semialdehyde (ACMS), which can cyclize non-enzymatically to quinolinic acid for the synthesis of a mononucleotide NAD precursor ^19,23,24^ by quinolinate phosphoribosyltransferase (QPRT). Combined deficiency of dietary Trp and vitamin B3 (niacin) causes systemic NAD depletion and pellagra. Alternatively, ACMS is decarboxylated by ACMSD to 2-aminomuconate-6-semialdehyde (AMS), which decays non-enzymatically to picolinic acid ^25^ unless oxidized to 2-aminomuconate (2-AM) by ALDH8A1 ^26^. Like preceding intermediates, 2-AM is chemically labile and undergoes spontaneous deamination to form 2-oxohex-3-enedioate (**Fig. 1a**).

Despite decades of biochemical investigation, the metabolic fate of 2-AM in humans and other eukaryotes remains unresolved ^27–29^. Early radiotracer studies indicated that Trp carbon enters the common 2-ketoadipate route, ultimately producing glutaryl-CoA and acetyl-CoA for mitochondrial metabolism ^20^. Consistently, an NAD(P)H-dependent activity capable of converting 2-AM into 2-ketoadipate was reported in cat liver extracts ^22^, although this assignment remained tentative owing to the substrate instability and the limitations of partially purified enzymes. By contrast, analysis of the KP gene cluster in the bacterium *Burkholderia cenocepacia* revealed a dedicated 2-aminomuconate deaminase (AMD) that efficiently converts 2-AM to 2-oxohex-3-enedioate ^30^. Whether an analogous or distinct enzymatic solution operates in eukaryotes has remained unknown.

## Results

### AMD homologs are involved in fungal KP

Because bacterial AMD belongs to the widespread IPR006175 family, we analysed this family across fungi and metazoa. Metazoans, including humans, typically encode a single member, RIDA (reactive imine deaminase), whereas fungi often harbour multiple homologues. A subset of fungal proteins clusters with validated bacterial AMDs (**Fig. 1b**), including a representative from *Aspergillus flavus*, selected for functional characterization. In various ascomycetes, these genes occur in compact genomic regions (5-10 kb) together with other components of the kynurenine pathway ^31^, supporting their assignment as fungal AMDs (**Fig. 1c**). Despite variation in local gene order, one architectural feature is conserved: HAAO and ACMSD are arranged head-to-head, with their 5′ ends separated by 550 ± 220 bp. Notably, all pathway genes contain introns except AMD.

To test whether the *A. flavus* and *H. sapiens* homologues catalyse the AMD reaction, we generated the unstable 2-AM substrate enzymatically from commercially available 3-HAA using three recombinant human enzymes (HAAO, ACMSD and ALDH8A1), whose activities were monitored by UV spectroscopy (**Fig. 1d** and **Supplementary Fig. S1a-c**). We subsequently isolated 2-AM by ion-exchange chromatography (**Supplementary Fig. S1d-f**) under mildly alkaline conditions (pH 9.5), with identity and purity confirmed by NMR (**Fig. 1e** and **Supplementary Fig. S1g**). Purified 2-AM underwent spontaneous deamination with a half-life of ∼160 min at pH 7.6 (**Fig. 1f** and **Supplementary Fig. S1h**). Addition of recombinant human RIDA did not measurably accelerate this reaction, whereas the *A. flavus* protein triggered rapid ammonia release and formation of a product with the same spectral features as the spontaneous decay product (**Fig. 1 g,h and Supplementary Fig. S1i and S2b,c**).

These results indicate that fungi, in contrast to metazoa, encode an AMD capable of accelerating the conversion of 2-AM to 2-oxohex-3-enedioate. Unlike human RIDA, AMD did not enhance the hydrolysis of 2-iminopropanate (**Supplementary Fig. S2d**), a reactive imine produced in threonine catabolism, consistent with a specialized role in the fungal KP.

### Coevolutionary screening identifies ASPDH as a candidate KP gene

KP genes exhibit coordinated patterns of gain and loss across eukaryotes, forming a coevolving network (**Fig. 2a**). We reasoned that this signal could reveal the missing component acting downstream of 2-AM. However, an untargeted computational screening, previously effective in other metabolic contexts ^32^, did not identify compelling candidates. We therefore implemented a targeted analysis, screening human genes for coevolution with known pathway components and controlling key variables affecting coevolutionary inference, including taxa selection, orthology assignment and coevolution metrics ^33–37^.

**Fig 2.**
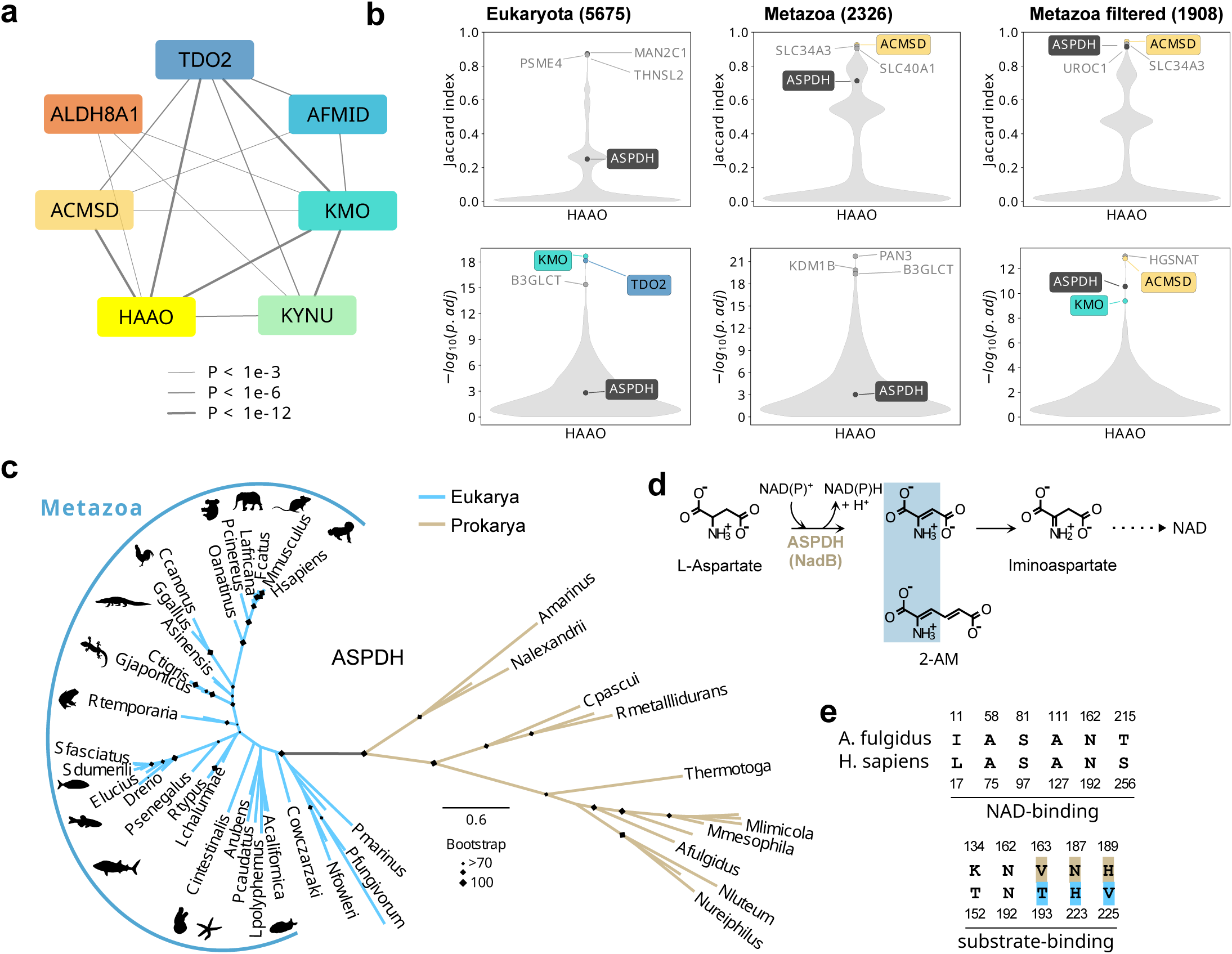
Identification of ASPDH as a candidate KP gene in humans. **a,** Coevolution network of human genes of tryptophan catabolism, arranged circularly according to pathway order; edges denote correlated gain/loss events across eukaryotes, weighted by statistical significance (cotr P values). **b,** Violin plots showing the distributions of Jaccard index (top) and cotr P values (bottom) obtained using HAAO as a bait in different data sets; the three best-scoring genes and the candidate ASPDH are indicated. Species sets and number of species are shown above each plot. **c,** Maximum-likelihood phylogeny of ASPDH across Bacteria, Archaea and Eukarya. **d,** Scheme of the known ASPDH reaction highlighting the chemical similarity of the involved compounds with 2-AM. **e,** Comparison of conserved NAD-binding and substrate-binding positions in ASPDH sequences from *H. sapiens* and *Archaeoglobus fulgidus*. Conserved differences between eukaryotic and prokaryotic sequences are highlighted.

Focusing the analysis on metazoan genomes, in light of the identification of a fungal-specific AMD, proved decisive (**Fig. 2b**). Under these conditions, a gene annotated as aspartate dehydrogenase (ASPDH), which was detected in the Eukaryota dataset only by a sensitive metric ^34^ with ACMSD as a bait (**Supplementary Fig S3a-f**), emerged as a coevolving partner of other in-pathway genes. Further refinement came from the recognition of false gene absence in several bird species ^38^, supported by tBLASTn recovery of avian ASPDH transcripts and by the high GC content of these sequences (**Supplementary Fig. S3g,h**). Excluding Aves strengthened the signal across different pathway genes and coevolution metrics (**Fig. 2b**). Under these conditions, reciprocal screening with ASPDH as bait identified KP genes as the most significant interacting partners (**Supplementary Fig. S3c,f**), further supporting ASPDH as a candidate.

Phylogenetic analysis showed a clear separation between prokaryotic and eukaryotic ASPDH sequences (**Fig. 2c**). Homologues are present in selected Bacteria and Archaea and, within Eukarya, are largely restricted to metazoans, being absent from fungi, plants and most unicellular eukaryotes. Notably, homologues are detected in the amoeboid protist *Capsaspora owczarzaki*, which branches close to metazoa, indicating an early presence in the holozoan lineage.

The functional annotation of ASPDH is based on its homology to prokaryotic L-aspartate dehydrogenase (nadB), involved in the formation of iminoaspartate ^39^ in the *de novo* pathway of NAD biosynthesis (**Fig. 2d**). However, metazoans use the KP for NAD biosynthesis (see **Fig. 1a**) and an enzyme able to oxidize L-aspartate was not detected in mammals tested for this activity ^40^. ASPDH belongs to a large oxidoreductase superfamily (CATH 3.30.360.10) with diverse substrate specificity. Notably, 2-AM displays chemical similarity to substrates and intermediates of the ASPDH reaction, in particular to a possible dehydro-aspartate intermediate (**Fig. 2d**). Residues involved in cofactor binding are largely conserved between prokaryotic and eukaryotic sequences, whereas substrate-binding residues show substantial divergence (**Fig. 2e**), suggesting conservation of oxidoreductase activity but altered substrate specificity.

### Human ASPDH functions as a NAD(P)H-dependent 2-AM reductase

Recombinant human ASPDH corresponding to the canonical isoform (283 aa) was expressed and purified, and eluted with an apparent molecular mass of ∼56 kDa, indicative of a dimer (**Supplementary Fig. S4a,b**). The protein did not catalyse the L-aspartate dehydrogenase reaction. Instead, in the presence of 2-AM, it exhibited strong NAD(P)H-dependent reductase activity, with ∼2-fold higher rates with NADPH than NADH (**Fig. 3a**). The reaction followed Michaelis-Menten kinetics with respect to 2-AM (**Supplementary Fig. S4c**), with a *K_m_* of 36.5 μM and a *k_cat_* of 2.9 s^−1^, corresponding to a catalytic efficiency (*k_cat_ / K_m_*) of ∼8.0 x 10^4^ M^−1^ s^−1^. NAD(P)H oxidation and 2-AM turnover occurred with a 1:1 stoichiometry (**Supplementary Fig. S4d**). Incorporation of ASPDH into the reaction cascade from 3-HAA prevented 2-AM accumulation (**Supplementary Fig. S4a**). However, rather than accelerating 2-AM deamination, ASPDH also prevented ammonium release (**Fig. 3b**), inconsistent with direct conversion to 2-ketoadipate.

**Fig 3.**
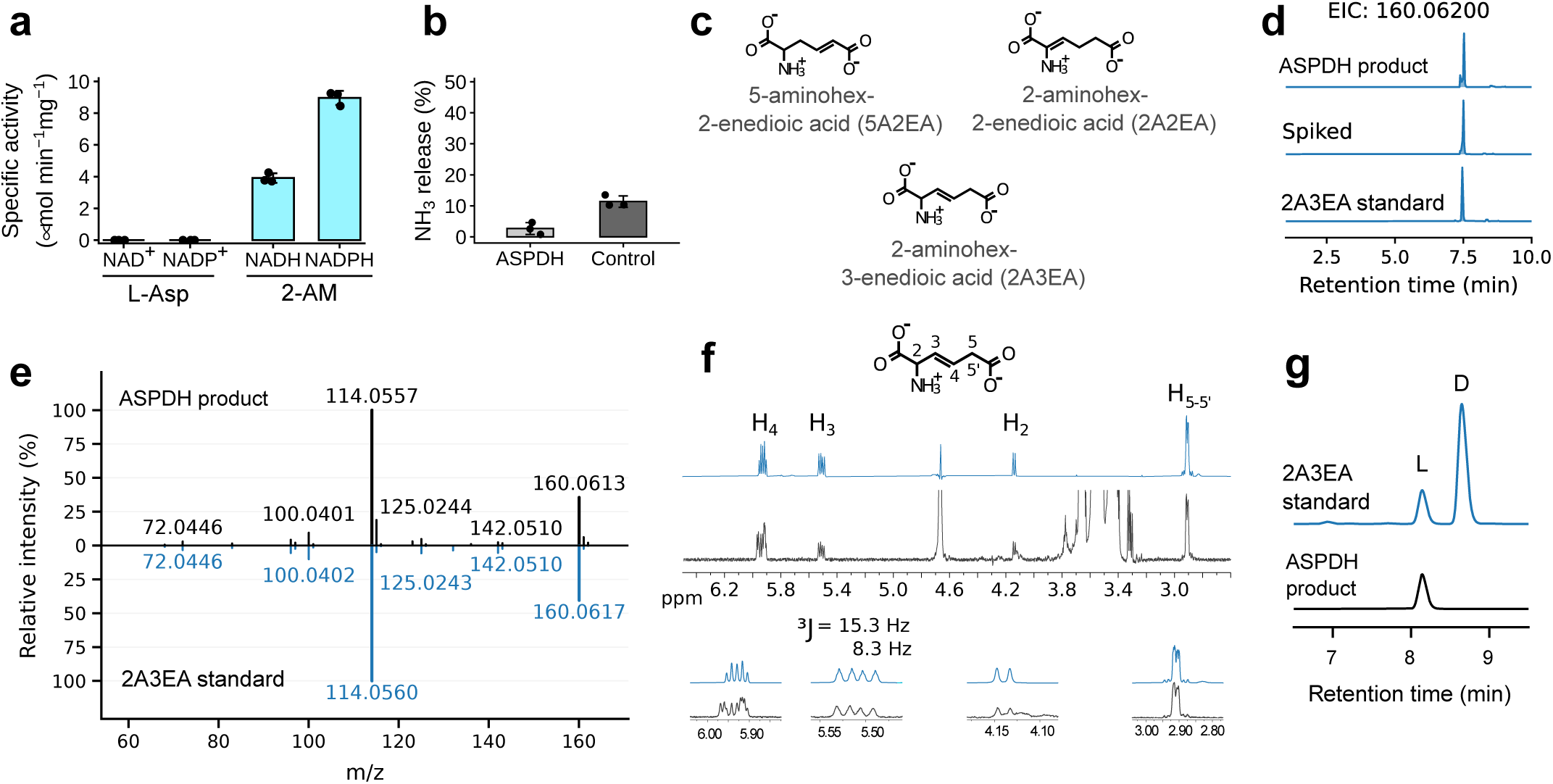
Human ASPDH catalyses the formation of 2A3EA in tryptophan catabolism. **a**, Specific activity of ASPDH towards L-Asp using NAD^+^/NADP^+^ and towards 2-AM using NADH/NADPH. **b**, Ammonium released from 2-AM in the presence of ASPDH and in a no-enzyme control. Points show individual measurements (n = 3); bars represent the mean ± SD. **c**, Chemical structures of the three putative ASPDH reaction products. **d**, Extracted ion chromatograms (EIC) from HILIC LC–MS for a chemically synthesised 2A3EA standard, the ASPDH reaction product, and a spiked sample. **e**, Comparison of mass fragmentation spectra between the ASPDH reaction product and the 2A3EA standard. **f**, ^1^H NMR spectrum (black) obtained after 5-minute addition of human ASPDH and NADH to 2-AM compared with the spectrum of 2A3EA standard (blue). Corresponding protons are indicated on the chemical structure above. **g**, Stacked chromatograms of Marfey-derivatized 2A3EA standard and ASPDH product analysed by reversed-phase HPLC (340nm), showing enantioseparation of derivatized 2A3EA.

Recent work identified ASPDH as a candidate binding protein for nicotinic acid adenine dinucleotide phosphate (NAADP) ^41^ though the physiological relevance of this interaction remained unclear. We found that NAADP inhibits AMR activity in a concentration-dependent manner across multiple NADH concentrations, consistent with competitive inhibition (*K_i_* = 930 nM; **Supplementary Fig. S4e**). The non-phosphorylated analogue NAAD did not inhibit activity under identical conditions (**Supplementary Fig. S4f**).

Together, these results establish ASPDH as the 2-AM reductase (AMR) of the animal KP, although the reaction product did not correspond to any previously recognized metabolite of the pathway.

### Human ASPDH/AMR produces 2-aminohex-3-enedioic acid

We next sought to define the outcome of the AMR reaction. In the absence of ammonium release and with stoichiometric consumption of NADH, the product is expected to retain a double bond, yielding either 2-aminohex-2-enedioic acid (2A2EA) or 5-aminohex-2-enedioic acid (5A2EA), depending on the site of hydride addition (**Fig. 3c**). Both compounds have a monoisotopic mass (159.053 Da) consistent with the product observed by mass spectrometry (**Fig. 3d,e** and **Supplementary Fig. S5a-c**). However, ^1^H NMR spectrum (**Fig. 3f**) showed signals inconsistent with those expected for these two isomers.

We therefore considered the possibility that the reaction involves rearrangement of the conjugated system, generating 2-aminohex-3-enedioic acid (2A3EA), in which the double bond is shifted to the C3–C4 position (**Fig. 3c**). This structure accounts for the observed spectroscopic features, and COSY analysis revealed a coupling topology fully consistent with 2A3EA (**Supplementary Fig. S5d**). This assignment was confirmed by comparison with a synthetic standard: the enzymatic product co-eluted with 2A3EA, displayed identical exact mass and MS/MS fragmentation profiles, and showed complete overlap in both one- and two-dimensional NMR spectra (**Fig. 3d–f** and **Supplementary Fig. S5**).

In both the enzymatic product and the reference compound, the vinylic protons exhibited a large vicinal coupling constant (³J ≈ 15.3 Hz), consistent with a *trans* (E) configuration of the double bond (**Fig. 3f**). Upon derivatization with a chiral reagent and reversed-phase chromatography, chemically synthesized 2A3EA resolved into two peaks, whereas the enzymatic product yielded a single peak (**Fig. 3g and Supplementary Fig. S6**), which was assigned the L (2S) configuration based on established elution orders ^42^ and comparison with enantiopure analogs. Together, these data identify (2S,E)-2A3EA as the product of the AMR-catalysed reaction. This unsaturated α-amino acid had previously been obtained as a racemic mixture by chemical synthesis ^43^.

### Mechanism of 2A3EA formation in tryptophan catabolism

To elucidate the mechanism of 2A3EA formation in the AMR reaction, we sought structural insight into the enzyme–substrate complex. Despite extensive efforts, diffracting crystals of recombinant AMR could not be obtained. However, a high-confidence ternary AMR–NADH–2-AM complex (pTM = 0.93, ipTM = 0.93), with both ligands stably positioned in the active site (pairwise ipTM ≈ 0.94), was predicted with AlphaFold3 (AF3) (**Fig. 4a and Supplementary Fig. S7**).

**Fig 4.**
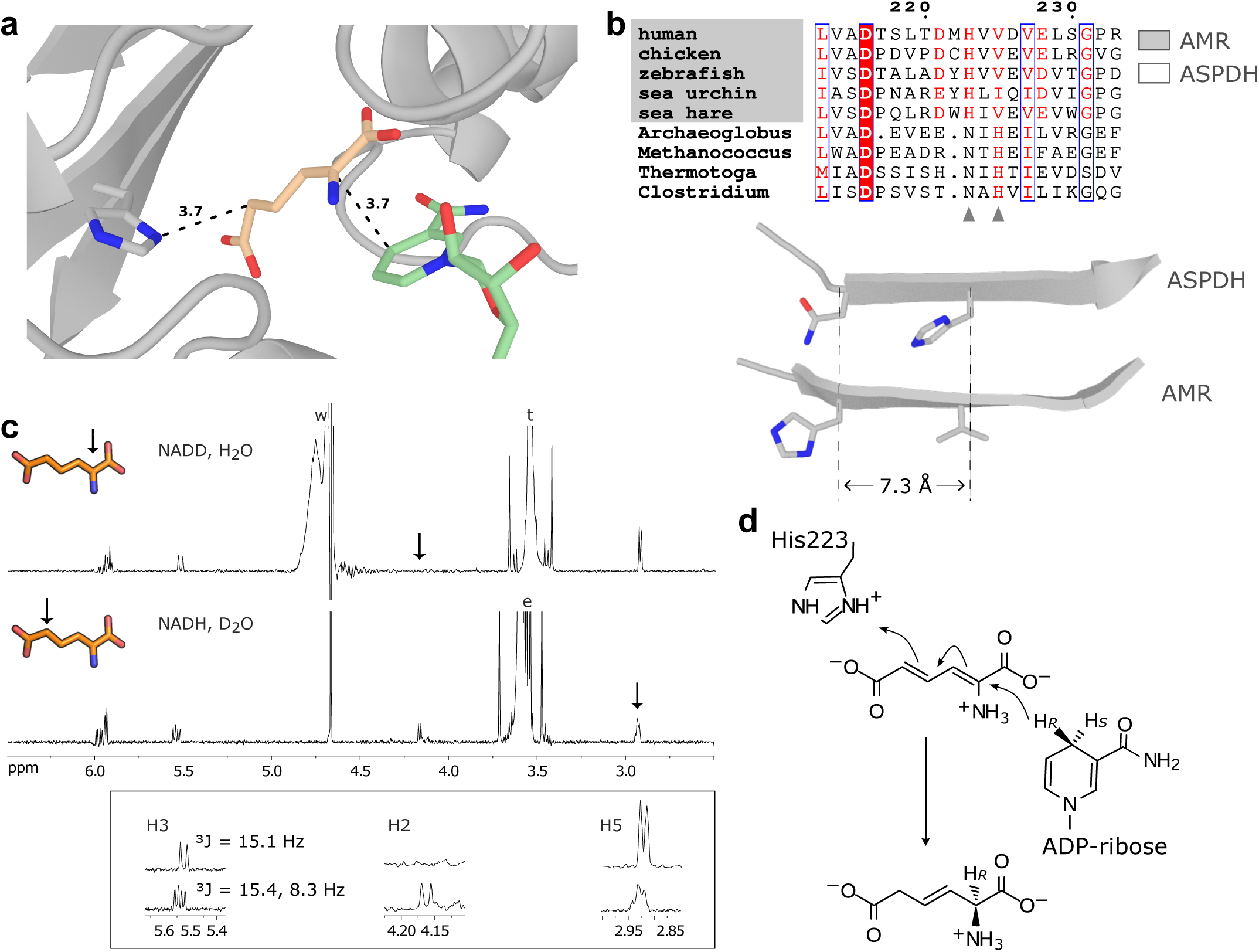
Structural and mechanistic basis of 2A3EA formation by AMR. **a**, AlphaFold3 prediction of the ternary AMR–NADH–2-AM complex, showing placement of 2-AM (wheat carbon) and nicotinamide ring (palegreen carbon) ligands within the active site. Distances (Å) between nicotinamide C4 and 2-AM C2, and between His223 and 2-AM C5, are indicated. **b**, Divergence of the catalytic histidine between ASPDH and AMR. Upper panel, excerpt of a sequence alignment from representative species, with the active-site histidine indicated by arrowheads. Lower panel, structural alignment showing a 7.3 Å shift of the His223 alpha carbon. **c**, Representative ¹H NMR traces of reactions conducted with NADD in H_2_O (top) and NADH in D_2_O (lower). Positions of deuterium incorporation, revealed by loss of ¹H signal, are indicated by arrows. Signals from H_2_O (w), Tris–HCl buffer (t), and undeuterated ethanol (e) used for NADH recycling are labeled. **d**, Proposed mechanism for 2A3EA formation, in which hydride transfer to C2 is coupled to reorganization of the conjugated π system and proton transfer at C5.

The geometry of the predicted ternary complex provides a coherent framework for the reaction sequence. In the model, the pro-R hydride of NADH is aligned for transfer to C2 of 2AM. The trajectory of hydride delivery approaches the si face of the 2-AM π-system, thereby specifying formation of the L configuration at the newly generated stereocenter. The model further predicts protonation at C5, enabled by the positioning of the substrate relative to a conserved active site histidine (**Fig. 4a,b)**.

These mechanistic features inferred from the AF3 model were fully supported by experimental evidence. When the reaction was monitored by ¹H NMR using [4R-²H]NADH, we observed loss of the H2 proton signal in the product, together with loss of the H3-H2 coupling (^3^J = 8.3 Hz), indicating hydride transfer from the pro-R position of NADH to C2 of 2-AM (**Fig. 4c, upper panel**). Conversely, when the reaction was performed with undeuterated NADH using enzyme preparations buffer-exchanged into D₂O, one of the two geminal proton signals at C5 of the product was selectively lost **(Fig. 4c, lower panel**), indicating site-specific proton transfer at C5 mediated by the enzyme.

These data support a mechanism in which hydride transfer to C2 is coupled to reorganization of the conjugated π system (**Fig. 4d**), driving migration of the double-bond to the β–γ position and enabling proton transfer at C5. This coupling accounts for the stereochemical and regioselective formation of (2S,E)-2A3EA as the direct product of the AMR reaction.

Interestingly, the histidine residue (His223) implicated in proton transfer at C5 in our model is among the residues that differ between prokaryotic ASPDH and eukaryotic AMR (**Fig. 2e** and **Fig. 4b**, **upper panel**). In AMR, His223 is positioned two residues upstream relative to a conserved His in ASPDH. Because the residue lies on a β-strand, this sequence displacement preserves the side-chain orientation while resulting in an approximately 7 Å translation of the His α-carbon (**Fig. 4b**, **lower panel**). This repositioning places His in proximity to C5 rather than C3 of the substrate, thereby enabling proton transfer at C5, in contrast to what is observed in ASPDH. This rearrangement provides a structural explanation for the altered regioselectivity of proton transfer and defines the molecular basis for the evolutionary adaptation of the AMR catalytic mechanism.

### Catalytically active AMR is expressed in liver and kidney

The human ASPDH locus encodes two protein isoforms that differ in length, domain architecture and tissue distribution (**Supplementary Fig. S8**). Isoform 1 (i1) corresponds to the full-length protein, whereas isoform 2 (i2) is a shorter variant generated from an alternative transcriptional start site and exon skipping. Transcriptomic data indicate that i1 is predominantly expressed in liver and kidney, consistent with tissues in which the KP is most active, whereas i2 is relatively enriched in the brain, including the cerebral cortex (**Supplementary Fig. S8a-d**).

The two isoforms differ markedly in their predicted structural organization: i1 contains both the N-terminal Rossmann fold required for nicotinamide cofactor binding and the C-terminal dehydrogenase domain, whereas i2 lacks the majority of the Rossmann domain together with the canonical NAD(P)-binding motif (**Supplementary Fig. S8e,f**). This truncation predicts loss of oxidoreductase activity in i2. Consistent with this prediction, recombinant i2 (**Supplementary Fig. S4g,h**) did not show detectable activity toward 2-AM under conditions in which i1 displayed high catalytic turnover (**Supplementary Fig. S8g**).

While the main isoform is annotated in most metazoan genomes, i2 is annotated only in humans and close primate relatives (Catarrhini). Comparative analysis of the genetic locus in 30 primate species reveals that i2 is potentially encoded only by Simiiformes (true monkeys) among mammals (**Supplementary Fig. S8h**). This pattern indicates that i2 represents a recent evolutionary innovation rather than an ancestral alternative state of the enzyme.

Together, these data define a clear functional partitioning within the human genetic locus: isoform 1 is the conserved, catalytically active AMR operating in peripheral metabolic tissues, whereas isoform 2 is a brain-enriched, catalytically inactive variant of recent evolutionary origin.

### Evolutionary history of the metazoan kynurenine pathway

We examined at the species-level detail the correlated gene-loss and retention patterns that enabled the identification of AMR. The analysis was conducted with a more accurate HMM-based orthology classification, including the known genes of the pathway (see **Fig. 1a**), and the newly identified AMD and AMR genes (**Supplementary Fig. S9a**).

At the kingdom level, pathway gene distribution highlights a strong enrichment of the KP in Opisthokonta (including fungi and metazoa), whereas plants retain only QPRT, which functions in the aspartate-derived NAD biosynthesis pathway. This analysis further supports the mutually exclusive presence of AMD and AMR in fungi and metazoa, respectively (**Supplementary Fig. S9b)**.

Within metazoa, AMR is detected from early-branching clades, such as Porifera (for example, *Amphimedon queenslandica*) and Cnidaria (for example, *Hydra vulgaris*), to derived deuterostome lineages (**Supplementary Fig. S9b,c**). The only pathway truncation observed in deuterostomes is the loss of QPRT in the Ostariophysi, a large teleost clade including zebrafish, goldfish and catfish, consistent with previous experimental evidence ^44,45^. By contrast, several correlated gene losses are observed in protostomes, with the systematic lack of the HAAO-AMR branch in insects and the independent loss of the whole pathway in flatworms (Platyhelminthes), with the exception of basal Rhabditophora, and in roundworms (Nematoda), with the exception of *C. elegans* and related Rhabditida. More taxonomically restricted truncations are observed in Cephalopoda within Mollusca and in Myxozoa within Cnidaria (**Supplementary Fig. S9c**).

Pathway truncation in insects likely reflects an evolutionary trade-off, whereby the 3-HK intermediate is diverted towards ommochrome biosynthesis. Among the 883 insect species analysed, only three encode AMR. These species, representing the family Sciaridae (Diptera), uniquely retain the upstream components of the pathway, which are organized in a conserved genomic cluster including AMR, HAAO, ACMSD, as well as homologs of ALDH8A1 and AFMID, which are also present in other insects. Sequence similarity of this locus supports its bona fide dipteran origin (**Supplementary Fig. S9d**). The nested position of Sciaridae within Insecta supports a secondary re-acquisition of the pathway following its ancestral loss. Consistently, Sciaridae have also regained QPRT at a distinct genomic locus (**Supplementary Fig. S9d)**, indicating a restored capacity to derive NAD from tryptophan.

Overall, this analysis supports a tight evolutionary and functional coupling between ASPDH/AMR and the other KP genes across metazoa. The coevolutionary signal is mostly provided by protostome lineages.

## Discussion

Our results reveal two distinct solutions for the late steps of tryptophan catabolism in eukaryotes. Fungi encode a bacterial-like 2-aminomuconate deaminase that accelerates the spontaneous decay of this unstable intermediate, whereas metazoans instead employ an NAD(P)H-dependent reductase, here identified as ASPDH/AMR, to generate (2S,E)-2-aminohex-3-enedioic acid. This mutually exclusive distribution defines an evolutionary bifurcation in tryptophan catabolism downstream of 2-aminomuconate.

The emergence of AMR illustrates how a conserved dehydrogenase scaffold can be repurposed to support a new metabolic function. Although homologous to prokaryotic aspartate dehydrogenases, the metazoan enzyme has diverged in substrate specificity and reaction outcome while retaining the redox cofactor machinery. A minimal structural change—repositioning of a catalytic histidine along a β-strand—appears key to redirecting proton transfer and enabling a coupled hydride transfer and double-bond migration. A related mechanism is observed in fatty acid metabolism, where 2,4-dienoyl-CoA reductases process conjugated intermediates through reduction coupled to double-bond migration, although in that case rearrangement of the double-bond system is mediated by the enoyl-CoA chemistry ^46^.

Recent studies on liver cancer cells have implicated ASPDH in lactate metabolism and immune regulation ^47^, but interpreted the protein as a canonical aspartate dehydrogenase. Our findings suggest that previously observed phenotypes may arise from modulation of tryptophan catabolism and cellular redox balance rather than direct aspartate oxidation.

The identification of (2S,E)-2-aminohex-3-enedioic acid, for which we propose the name *latensine* (from Latin *latens*, hidden, reflecting its long-standing absence from metabolic maps), expands the known repertoire of endogenous metabolites and reveals a previously unrecognized metazoan branch of the kynurenine pathway. As an unsaturated α-amino acid not represented in current biological databases, latensine highlights how even well-studied metabolic routes can harbour undetected intermediates and how the growing availability of evolutionary diverse genomes can help uncover them.

Latensine is compatible with the proposed 2-ketoadipate/glutaryl-CoA terminal route of tryptophan catabolism, but implies the existence of additional metabolic steps and possibly further intermediates. The absence of additional coevolving partners suggests that subsequent steps may involve enzymes with broad substrate specificity or complex evolutionary histories that escape detection by current approaches. From a chemical standpoint, transformations such as transamination, hydration or deamination appear plausible. However, the identification of a stable product downstream of a sequence of highly labile intermediates (see **Fig. 1a**) provides a tractable entry point for pathway reconstruction and future discoveries.

## Supplementary Figure Legends

**Fig S1. Enzymatic synthesis and purification of 2AM.**

**a–c,** IMAC elution profiles (left) and SDS–PAGE gels (middle) showing purification of (a) HAAO, (b) ACMSD and (c) ALDH8A1. Lanes: marker (M), non-induced (NI), induced (I), supernatant (S), flow-through (FT), and numbered fractions, with matching fraction numbers between profiles and gels. UV–visible spectral changes (right) monitor substrate conversion by the purified enzymes. For HAAO, the first recorded spectrum (yellow) corresponds to 3-HAA prior to enzyme addition; subsequent spectra were acquired starting at 1 min after HAAO addition and every 30 s thereafter. For ACMSD and ALDH8A1 assays, upstream enzymes were included to generate substrates *in situ* (HAAO for ACMSD; HAAO and ACMSD for ALDH8A1), and reactions were initiated by addition of 3-HAA; spectra acquired every 30 s. **d**, UV–visible spectral changes during the one-pot enzymatic reaction leading to 2-AM formation. The reaction was initiated by addition of 3-HAA; spectra were acquired every 1 min, with the final spectrum recorded 2 min after the previous time point. **e**, Anion-exchange chromatography profile of the 2-AM formation reaction mixture. Absorbance at 260 (pinkish red) and 328 (cyan) nm is plotted versus elution volume. Peaks were assigned based on their UV–visible spectral properties. **f**, UV-visible absorption spectrum of purified 2-AM. **g**, COSY NMR spectrum of purified 2-AM, showing peaks at 7.15 (H4), 5.9 (H3), and 5.8 (H5) ppm. Proton assignments correspond to the chemical structure. **h**, NADH oxidation was monitored at 340 nm in the presence of glutamate dehydrogenase and α-ketoglutarate after addition of 2-AM, with time zero defined at 80 s. The ratio [NADH oxidized]/[2-AM] approaches unity consistent with a 1:1 stoichiometry between NH₄⁺ release and NADH oxidation. **i**, UV–visible absorption spectrum of 2-AM after overnight incubation.

**Fig S2. Expression and functional characterization of AMD eukaryotic homologs.**

**a**, IMAC elution profile (left) and SDS–PAGE gel (right) showing purification of RIDA. Lanes: marker (M), non-induced (NI), induced (I), supernatant (S), flow-through (FT), pellet (P) and numbered fractions, with matching fraction numbers between profile and gel.

**b**, SDS-PAGE gel (left) showing *A. flavus* AMD. Lanes: marker (M) and enzyme in duplicate (AMD). UV–visible spectral changes (right) following addition of *A. flavus* AMD to purified 2-AM in with an excess enzyme (0.5 μM), showing an enzyme-independent spectral transition. The first spectrum (dark blue) corresponds to 2-AM before enzyme addition; subsequent spectra were acquired starting at 5 s after AMD addition and every 30 s thereafter. **c**. COSY NMR spectrum of 2-oxohex-3-enedioate, obtained from purified 2-AM in presence of *A. flavus* AMD, showing peaks at 3.16 (H5-H5’), 6.95 (H4), and 6.20 (H3) ppm. Proton assignments are shown in the corresponding chemical structure. **d**, Schematic reaction (left) of RIDA family activity coupled with L-threonine dehydratase (Ilva) to produce 2-ketobutyrate. µM min-1 of NADH consumed (right) from RIDA activity in a three-enzyme coupled assay with GDH, Ilva and the enzymes from *A. flavus* (AMD) and *H. sapiens* (RIDA), and in a no-enzyme control.

Points show individual measurements (n = 3); bars indicate the mean ± SD.

**Fig S3. Coevolutionary screening for KP genes.**

**a-f**, Violin plots showing the distributions of cotr P values (**a–c**) and Jaccard indices (**d–f**) obtained using ACMSD (**a,d**), ALDH8A1 (**b,e**), and ASPDH (**c,f**) as bait genes in eukaryotic (left; n species = 5,675, n genes = 77,042,201), metazoan (middle; n species = 2,326, n genes = 38,031,849), and metazoan excluding birds (right; n species = 1,908, n genes = 32,131,031) datasets. The three top genes, together with HAAO, ACMSD, ALDH8A1, and ASPDH, are indicated. Note that the inclusion of ALDH8A1 within a large orthogroup containing multiple aldehyde dehydrogenases obscures its coevolutionary relationships with other KP genes. **g**, Phylogenetic distribution of kynurenine pathway genes across selected Metazoa, with emphasis on Aves (grey shading). The heatmap indicates the presence (gold) or absence (white) of pathway enzymes across species ordered according to the rooted NCBI taxonomy tree. **h**, Table of selected avian species lacking annotated ASPDH protein sequences, for which corresponding transcript sequences were identified by tblastn using the *Gallus gallus* ASPDH protein (UniProt: A0A8V0X7L2_CHICK) as query. Columns report organism names, mRNA accession numbers (EST and TSA datasets), sequence identity (%), E-values relative to the *Gallus gallus* ASPDH protein, and mRNA G+C content.

**Fig S4. Human ASPDH functions as 2-aminomuconate reductase (AMR).**

**a,** IMAC elution profiles (left), SDS–PAGE analysis (middle) and UV–visible spectral changes (right) for ASPDH canonical isoform (Uniprot A6ND91). Lanes: marker (M), non-induced (NI), induced (I), supernatant (S), flow-through (FT), and numbered fractions, with matching fraction numbers between chromatograms and gels. Upstream enzymes (HAAO, ACMSD and ALDH8A1) were included to generate substrates *in situ*, and reactions were initiated by addition of 3-HAA; spectra were acquired every 30 s. The decrease in the 2-AM signal is consistent with substrate turnover by ASPDH. **b,** SEC elution profile of ASPDH, with an estimated molecular mass of ∼56 kDa. MW standards are indicated (BSA dimer, 132 kDa; BSA monomer, 66 kDa; trypsinogen, 24 kDa). **c,** Michaelis–Menten kinetics of ASPDH with 2-AM as substrate. Data points represent mean values (n = 3); error bars indicate ± SD; line indicates the best fit to the Michaelis–Menten equation. **d,** Top, NADPH oxidation at 340 nm was monitored in the presence of 2-AM following addition of ASPDH (∼70 s). The [NADPH oxidized]/[2-AM] ratio approaches unity consistent with a 1:1 stoichiometry between NADPH oxidation and 2-AM reduction. Bottom, stoichiometry of 2-AM reduction relative to cofactor oxidation in ASPDH reactions containing NADH or NADPH. Bars indicate mean ± SD and dots indicate individual replicates (n = 3). **e,** Dependence of the initial rate of NADH oxidation and 2-AM reduction by ASPDH on increasing NAADP concentrations, showing inhibition of enzyme activity. Data points represent individual measurements and were fitted by global analysis using a competitive inhibition model. **f,** Time course of ASPDH activity (0.5 µM) at 340 nm using purified 2-AM in the presence of NADH (150 µM) and either NAAD (150 µM, top left) or NAADP (150 µM, bottom left). NAADP inhibited the reaction, whereas no inhibition was observed with NAAD. Right, schematic representation of the connection between tryptophan catabolism and NAD biosynthesis, highlighting NAADP, a NAD-derived metabolite, as an inhibitor of the ASPDH-mediated conversion of 2-AM to 2A3EA. **g,** IMAC elution profiles (left), SDS–PAGE analysis (middle) and UV–visible spectral changes (right) for ASPDH shorter isoform (Uniprot A6ND91-2), which eluted during the wash steps; the asterisk (*) indicates purified protein obtained after a second IMAC step. Under the same assay conditions as in (a), the 2-AM peak persisted, indicating lack of catalytic activity. **h,** SEC elution profile of ASPDH (shorter isoform), with an estimated molecular mass of ∼69 kDa. MW standards are as in panel b.

**Fig S5. Spectroscopic evidence for 2A3EA as the ASPDH reaction product.**

**a,** Detail of the extracted ion chromatogram (EIC) in the analyte elution window of MSe acquisition; expanded view of the retention time region 7.0–8.0 min showing the EIC at *m/z* 160.061 ± 20 mDa ([M+H]⁺, C₆H₁₀NO₄⁺) for three sample conditions. The spiked sample was prepared by addition of the reference standard to the sample matrix prior to analysis. **b,** MS/MS fragmentation map of 2A3EA. Positive-ion electrospray ionisation MS/MS spectrum was obtained using an optimised collision energy ramp. Relative intensities are normalised to the base peak (*m/z* 114.056, [M+H−H₂O−CO]⁺, 100%). Coloured arrows indicate fragmentation pathways connecting precursor and product ions: dark green, −46.00 Da (−H₂O, −CO); red, −60.02 Da (−C₂H₄O₂); orange, −18.01 Da (−H₂O); teal, −17.03 Da (−NH₃); blue, −27.99 Da (−CO); light green, −14.01 Da (−CH₂); yellow, −35.03 Da (−NH₃ and −H₂O); purple, −59.03 Da (−NH₃ and −COCH₂); brown, −42.01 Da (−COCH₂). Proposed ionic structures are shown above the precursor and the major fragment ions. **c**, List of MS/MS fragments.

Theoretical fragment *m/z* values, proposed neutral losses, mass errors relative to the closest centroid peak in the experimental spectrum (Δ ppm), and relative intensities are reported. **d,** Overlay of the COSY NMR spectra of the enzymatic product (black) and the 2A3EA standard (blue), with proton-coupling assignments.

**Fig S6. Chromatographic analysis of 2A3EA enantiomers after derivatization with Marfey reagent.**

**a**, Stacked chromatograms of Marfey-derivatized standards and samples analysed by reversed-phase HPLC (340 nm). **b**, Expanded view (5.5–13.5 min). α-Aminoadipic acid (AA) (racemic, D and L) was used as a standard to validate chromatographic separation of diastereomers. The AA racemic standard resolves into two peaks that co-elute with the respective enantiopure standards. A chemically synthesized 2A3EA sample yields two peaks, whereas the enzymatic product (enz-2A3EA) yields a single peak. **c**, Overlay of 2A3EA and enz-2A3EA chromatograms on a common y scale highlighting the elution region of the derivatized product. Peaks assigned to L-2A3EA, D-2A3EA and Tris–HCl (Trizma) are indicated. Based on the established elution order of Marfey derivatives in reversed-phase chromatography, the enzymatic product is assigned the L configuration.

**Fig S7. AlphaFold3 protein-ligand complexes and confidence metrics.**

**a-b**, AlphaFold3 predictions of the monomeric and dimeric forms of AMR bound to NADH and 2AM, colored by per-residue (protein) or per-atom (ligands) pLDDT. Panels below display the corresponding predicted aligned error (PAE) plots of the reference model (seed1), indicating confidence in inter-residue distances. **c-d**, Violin plots of ipTM, pTM, mean PAE, and ligand RMSD across all models from 100 independent seed runs. Each violin shows the full distribution; central markers indicate the mean. The red dot indicates the reference model (seed1). **e-h**, Same metrics grouped by AlphaFold3 ranking score within each run (1 = highest ranking_score). The red dot indicates the reference model (seed1).

**Fig S8. Isoform expression and characterization of the human ASPDH locus.**

**a**, Tissue expression profile of ASPDH based on the Human Protein Atlas (HPA) consensus nTPM values. **b**, Gene-wise normalized heatmap of tryptophan catabolism genes across the ten tissues with highest average expression. **c**, NCBI view of the ASPDH locus showing curated isoforms 1 (I1) and 2 (I2) with RNA-seq tracks for cerebral cortex, kidney and liver. **d**, Kallisto-based quantification of I1 and I2 transcripts in brain, kidney and liver (ENA: PRJNA764684). **e**, Sequence alignment of I1 and I2 highlighting the boundaries of the NAD(P)-binding and aspartate dehydrogenase (ASP_DH) domains (InterPro). **f**, Superimposed AlphaFold3-predicted structures of ASPDH (grey) and the hypothetical I2 (coloured by per-residue pLDDT scores). **g**, Time course at 326 nm with purified 2-AM in the presence of NADH (60 µM) demonstrating lack of activity after addition of I2 (2 µM); addition of I1 (1 µM) as a control results in rapid signal decrease. **h**, TimeTree mammalian phylogeny showing species potentially encoding I1 and I2, inferred from comparative analysis of the genomic locus in the UCSC Genome Browser.

**Fig S9. Evolution of tryptophan catabolism via the kynurenine pathway in eukaryotes.**

**a,** Distribution heatmap of KP genes across 5,570 eukaryotic species. Gene presence (rows) was assessed across organisms (columns) using profile HMMs; colours indicate normalized hmmsearch scores. Taxonomic slices corresponding to Fungi, Metazoa, Sar (S), and Viridiplantae (Vir) are indicated. **b,** Fraction of species (Aves excluded) encoding each pathway enzyme across eukaryotic kingdoms (top) and metazoan phyla (bottom). Dot size and colour indicate the proportion of genomes with an assigned gene (threshold = 0.3; group sizes in parentheses). **c,** Phylogenetic distribution of pathway genes across selected eukaryotes with emphasis on Metazoa. The heatmap indicates presence (orange) or absence (white) of KP enzymes across a time-calibrated species tree inferred by maximum likelihood ^34^ and dated using TimeTree ^48^. Major metazoan clades showing lineage-specific gene loss or retention are indicated. The tree is ladderized, with some nodes rotated to highlight independent loss and retention events. **d,** Microsynteny of the ASPDH locus in Sciaridae (Diptera), showing conserved genomic association with other KP genes. The table below reports the percentage identity and taxonomic origin of the best BLAST hit excluding Sciaridae.

Gene abbreviations: TDO2, tryptophan 2,3-dioxygenase; IDO1/2, indoleamine 2,3-dioxygenase; AFMID/KFA, arylformamidase/kynurenine-forming amidohydrolase; KMO, kynurenine 3-monooxygenase; KYNU, kynureninase; HAAO, 3-hydroxyanthranilate 3,4-dioxygenase; ACMSD, aminocarboxymuconate semialdehyde decarboxylase; AL8A1 (ALDH8A1), aldehyde dehydrogenase 8 family member A1; ASPDH, aspartate dehydrogenase; QPRT, quinolinate phosphoribosyltransferase; AMD, 2-aminomuconate deaminase.

## Methods

### Reagents

NADH (N8129), NAD (N7004), NADPH (481973), NADP (10128031001), NAAD (N4256), NAADP (N5655), 2-oxoadipic acid (2-OA; 75447), Glutamate dehydrogenase (GDH; G2626), α-ketoglutaric acid (K1128), Glucose-6-Phosphate Dehydrogenase (G6PDH; G8529), D-Glucose-6-Phosphate (G6P; G7250), Alcohol Dehydrogenase (ADH; A7011) were purchased from Sigma Aldrich. 3-Hydroxyanthranilic acid (3-HAA; X47470) was purchased from Organic Manchester. 2-Aminohex-3-enedioic acid (2A3EA) was purchased from Enamine (Kyiv, Ukraine). rac-ɑ-Aminoadipic acid (98%, Alfa Aesar, CAS 542-32-5), L-ɑ-aminoadipic acid (≥97.0%, TCI, CAS 1118-90-7), D-ɑ-aminoadipic acid (≥95.0%, TCI, CAS 7620-28-2), (S)-2-((5-fluoro-2,4-dinitrophenyl)amino)propanamide or 1 - fluoro-2,4-dinitrophenyl-5-L-alanine amide (L-FDAA or Marfey’s reagent ≥98.0%, TCI, CAS 95713-52-3), HPLC grade solvents (H_2_O, CH_3_CN) and analytical grade acetone were from VWR International S.r.l (Milan, Italy). Ethanol-d6 was purchased from Eurisotop (D114FD). NADH, NAD, NADPH and NADP stock solutions (15 mM) in water, and 2-OA, α-ketoglutaric acid, and G6P stock solutions (50 mM) in water were flash frozen in liquid nitrogen and stored at −80 °C. G6PDH was resuspended in 5 mM Glycine, pH 8, and stored at 4 °C.

### Cloning, expression and purification

The coding sequences of HAAO (NM_012205.2), ACMSD (NM_138326.2), ALDH8A1 (NM_022568.3), ASPDH - isoform 1 (NM_001114598.2), ASPDH - isoform 2 (NM_001024656.3), RIDA (NM_005836.2) were synthesized by Genscript (USA Inc.) into pET-28a(+) expression vector. The constructs were designed to add an N-terminal His-tag to the CDSs, with a cleavage site for either Thrombin or TEV protease (TEV site present only in ASPDH - isoform 2 construct). All plasmids were verified by DNA sequencing and transformed into E. coli BL21-Codon Plus.

Recombinant HAAO, ACMSD, ALDH8A1, ASPDH - isoform 1, ASPDH - isoform 2, and RIDA were expressed in auto-induction medium (LB supplemented with 0.05% glucose and 0.2% lactose) at 20 °C for 16 hours. Cells were harvested by centrifugation (20 min, 5000 × g, 4 °C), and resuspended in 40 mL lysis buffer (50 mM Tris-HCl pH 7.6, 300 mM NaCl). After lysis by sonication on ice (15 min of 1 s pulse at 35% intensity / 1 s pause), the soluble fraction was recovered by centrifugation (30 min, 20000 × g, 4 °C). Each protein was purified using a 5 mL His-trap Ni-NTA column (GE Healthcare) equilibrated in 50 mM Tris-HCl pH 7.6, with 300 mM NaCl, and eluted with an imidazole gradient (10-500 mM) on Äkta FPLC (GE Healthcare). The purified protein was buffer exchanged with PD-10 columns (GE Healthcare) into 100 mM potassium phosphate buffer pH 7.6, containing 150 mM NaCl. The final protein concentration was estimated by absorbance at 280 nm, using molar extinction coefficients computed with ProtParam (46535 M^−1^ cm^−1^ HAAO, 56295 M^−1^ cm^−1^ ACMSD, 65275 M^−1^ cm^−1^ ALDH8A1, 30730 M^−1^ cm^−1^ ASPDH - isoform 1, 8730 M^−1^ cm^−1^ ASPDH - isoform 2, 7450 M^−1^ cm^−1^ RIDA; https://web.expasy.org/protparam/). Aliquots were snap-frozen in liquid nitrogen and stored at −80 °C. AMD (XM_041287159.1) was synthesized by Genscript (USA Inc.) as a recombinant protein. The final protein concentration was estimated by absorbance at 280 nm using the molar extinction coefficient computed with ProtParam (30605 M^−1^ cm^−1^ for AMD).

The protein was aliquoted, snap-frozen in liquid nitrogen, and stored at −80 °C. Size-exclusion chromatography (SEC) was performed on a Superdex 200 Increase column (GE Healthcare) equilibrated in 100 mM potassium phosphate buffer pH 7.6, containing 150 mM NaCl, using an Äkta FPLC system (GE Healthcare).

### 2-AM preparation

The mixture for 2-AM preparation consisted of 100 mM potassium phosphate, pH 7.6, containing 1 mM 3-HAA, 2.5 mM NAD, 6 μM HAAO, 18 μM ACMSD, 24 μM ALDH8A1 in a final volume of 3.6 ml. The reaction was initiated by the addition of 3-HAA and followed spectrophotometrically. Over 160 minutes, the 330 nm absorbance peaked and then plateaued, confirming that the reaction had finished. 2-AM was subsequently purified using a modified version of a previously described protocol ^49^. Briefly, the reaction mixture was diluted to a final volume of 10.8 mL using 50 mM Tris-HCl (pH 9.5). To remove proteins, the mixture was filtered through a vivaspin tube (10 kDa cutoff; Amicon, Beverly, MA, USA). The filtrate was loaded onto a 5 mL HiTrap Q (MonoQ) ion-exchange column (GE Healthcare), pre-equilibrated with 10 mM Tris-HCl (pH 9.5). The column was washed with the same buffer until baseline absorbance was reached, then eluted using a 0–0.3 M NaCl linear gradient over 40 mL at a flow rate of 2 mL/min. Fractions of 0.5 mL were collected; 2-AM eluted in fractions 18–28. The concentration of 2-AM in each fraction was determined spectrophotometrically at 328 nm using ε328 = 16500 M⁻¹cm⁻¹. A total of 0.3 mg of 2-AM was obtained from 0.5 mg of 3-HAA. Aliquots were snap-frozen in liquid nitrogen and stored at –80 °C. To verify that purified 2-AM contained bound ammonium, an aliquot (80 µM) was incubated overnight in KP buffer (pH 7.6) containing NADH (200 µM), α-ketoglutarate (500 µM), and glutamate dehydrogenase (GDH, 2 U). Prior to the assay, GDH was purified using Vivaspin filtration to remove any residual ammonium contamination, ensuring that any ammonium detected originated from 2-AM.

### Activity assays

Catalytic activity measurements were carried out with a Jasco V-750 spectrophotometer equipped with a temperature control system and using a quartz cuvette of 1 cm path length. Assays were performed in 100 mM potassium phosphate buffer, pH 7.6, at 25 °C. HAAO activity was detected by the increase in absorbance at 360 nm due to the oxidative ring opening of 3-hydroxyanthranilate (3-HAA) to 2-amino-3-carboxymuconate semialdehyde (ACMS). Due to rapid kinetics, low enzyme concentrations (0.05 µM HAAO) were used to resolve the spectral shift from substrate (λmax 314 nm) to product (λmax 360 nm); under these conditions, the maximum product concentration was not reached due to spontaneous decay (ACMS half-life 36 min). ACMSD activity was detected by the decrease in absorbance at 360 nm as ACMS was converted to 2-aminomuconate 6-semialdehyde (AMS). ALDH8A1 activity was detected by the increase in absorbance at 330 nm due to the oxidation of AMS to 2-aminomuconate (2-AM) in the presence of NAD^+^. *A. flavus* AMD activity was measured using purified 2-AM, and activity was monitored by the decrease in absorbance at 326 nm and the increase in absorbance at 236 nm.

The GDH assay consisted of 50 mM potassium phosphate buffer (pH 7.6), 0.8 mM NH4Cl, 0.25 mM NADH, 0.5 mM α-ketoglutarate, and 2 U GDH. NADH oxidation was monitored at 340 nm.

Ammonium release from deaminase activity was quantified using a GDH-coupled assay consisting of 50 mM potassium phosphate buffer (pH 7.6), 0.15 mM NADH, 50 μM 2-AM, 0.5 mM α-ketoglutarate, 2 U GDH, and 0.5 µM *A. flavus* AMD or human RIDA. Buffer, 2-AM, α-ketoglutarate, and GDH were pre-incubated for 10 s, followed by addition of NADH. After 20 s, the reaction was initiated by addition of AMD or RIDA. Reaction progress was monitored at 340 nm.

RIDA family activity was measured using a three-enzyme coupled assay consisting of L-threonine dehydratase (IlvA), AMD or RIDA, and GDH. IlvA activity was measured using an IlvA–GDH coupled assay containing 50 mM potassium phosphate buffer (pH 7.6), 0.25 mM NADH, 10 mM L-threonine, 1 mM dithiothreitol (DTT), 0.5 mM α-ketoglutarate, 2 U GDH, and 1 µM IlvA. Buffer, L-threonine, DTT, α-ketoglutarate, NADH, and GDH were pre-incubated for 10 s prior to addition of IlvA. Reaction progress was monitored at 340 nm. The three-enzyme coupled assay consisted of 50 mM potassium phosphate buffer (pH 7.6), 0.25 mM NADH, 10 mM L-threonine, 1 mM DTT, 0.5 mM α-ketoglutarate, 2 U GDH, 1 µM IlvA, and 1 µM AMD or RIDA. Buffer, L-threonine, DTT, α-ketoglutarate, AMD or RIDA, NADH, and GDH were pre-incubated for 10 s prior to initiation by addition of IlvA. Reaction progress was monitored at 340 nm.

ASPDH activity was measured by the decrease in absorbance at 326 nm due to the reduction of 2-AM in the presence of NADH or NADPH. ASPDH activity was also tested using L-Asp as a substrate by monitoring absorbance changes at 340 nm. The reaction mixture consisted of 50 mM potassium phosphate buffer, pH 7.6, 60 µM L-Asp, 0.25 mM NAD+ or NADP+, and 0.05 µM ASPDH. The kinetic parameters of ASPDH were measured at different concentrations (4-80 μM) of 2-AM, in the presence of 0.25 mM NADH. Data of single wavelength (326 nm) kinetics were fitted to the Michaelis-Menten equation using the R package “drc”. The GDH assay consisted of 50 mM potassium phosphate, pH 7.6, 0.8 mM NH_4_Cl, 0.25 mM NADH, 0.5 mM α-ketoglutaric acid, and 1.2 U GDH.

The ASPDH-GDH coupled assay consisted of 50 mM potassium phosphate, pH 7.6, 0.25 mM NADH, 25 μM 2-AM, 0.5 mM α-ketoglutaric acid, 1 μM ASPDH and 1.2 U GDH. For the ASPDH reaction, potassium phosphate, pH 7.6, NADH, 2-AM, α- ketoglutaric acid, and ASPDH were combined and the reaction was incubated for 3 minutes. Then GDH was added and the reaction progression was followed at 340 nm absorbance.

NAADP inhibition kinetics were measured by the decrease in absorbance at 340 nm in the presence of varying NADH concentrations (30–300 μM) and NAADP (1–15 μM) in 50 mM potassium phosphate buffer, pH 7.6. Reaction rates were converted to μM min^−1^ using the extinction coefficients of 2-AM and NADH at 340 nm. Data were globally fitted to a competitive inhibition model using nonlinear least-squares regression implemented in the Python package “lmfit”.

The final product of AMD activity was analysed by ^1^H NMR spectroscopy. The reaction mixture consisted of 10 mM Tris-HCl, 357 µM 2-AM, 50 nM AMD, and 10% D_2_O in a final volume of 600 µL. The pH was adjusted to 7.78 using diluted HCl. ^1^H NMR spectra were acquired on a JEOL 600 MHz spectrometer approximately 30 min after enzyme addition. A COSY analysis was also performed on the same mixture.

### Isotope labelling

To investigate the reaction mechanism of AMR, reactions were analysed by ¹H NMR spectroscopy. Purified 2-AM was lyophilised (SP Scientific VirTis BenchTop Pro BTP-8ZC) prior to reaction setup. To minimise interference from NADH resonances in the ¹H NMR spectra, a low concentration of NADH was used and continuously regenerated using alcohol dehydrogenase (ADH) in the presence of ethanol. [4R-²H]NADH was prepared according to a described method ^50^ in a reaction mixture containing 100 mM NH₄HCO₃ buffer (pH 8.0), 0.16 mg alcohol dehydrogenase (ADH), 16 mM NAD, and 0.45 µL C₂D₅OD. After 15 min, the reaction mixture was diluted fivefold, and ADH was removed by ultrafiltration using a Vivaspin centrifugal filter unit (10 kDa molecular weight cutoff; Sartorius). The filtrate was loaded onto a 5 mL HiTrap Q ion-exchange column (GE Healthcare) pre-equilibrated with Milli-Q water. The column was washed with Milli-Q water until baseline absorbance was reached, and bound nucleotides were eluted with a 0–0.2 M NH₄HCO₃ linear gradient over 42 mL at a flow rate of 2.5 mL min⁻¹. Fractions (1 mL) were collected, and [4R-²H]NADH eluted in fractions 41–51. The concentration of [4R-²H]NADH in each fraction was determined spectrophotometrically at 340 nm using ε₃₄₀ = 6,220 M⁻¹ cm⁻¹.

In the first reaction, the mixture contained 0.35 mM 2-AM, 70 µM [4R-²H]NADH, 7 µM AMR, 35 mM ethanol-d6, 7U ADH, 50 mM potassium phosphate buffer (pH 7.6) and 10% D_2_O in a final volume of 600 µL. In the second reaction all components were prepared in a deuterated buffer and AMR preparations were buffer-exchanged into D₂O. The mixture contained 0.35 mM 2-AM, 70 µM NADH, 7 µM AMR, 4.5 mM ethanol, 7U ADH and 50 mM potassium phosphate buffer (pH 7.6) in a final volume of 600 µL. ¹H NMR spectra were acquired on a JEOL 600 MHz spectrometer ∼10 min after the enzyme addition; spectral changes were analysed with MestreNova V. 16.0.0-39276 to assess deuterium incorporation and infer mechanistic features of the reaction.

### LC-MS analysis of AMR reaction products

Chromatographic separation was performed on an Acquity UPLC I-Class Plus system (Waters, Milford, MA, USA) equipped with an Atlantis Premier BEH Z-HILIC column (1.7 µm, 2.1 × 100 mm; Waters). Mobile phase A consisted of water with 1 mM ammonium formate adjusted to pH 9.70 with 10% ammonium hydroxide; mobile phase B was acetonitrile.

Gradient elution was applied as follows: 0–2 min, 95% B; 2–7 min, 95–70% B; 7-8min, 70-60% B; 8–10 min, 70–10% B; 10–11 min, 10% B; 11–12 min, 10–30% B; 12–15 min, 30–95% B; flow rate 400 µL/min; column temperature 40 °C; injection volume 2.0 µL. Samples were maintained at 4 °C.

High-resolution mass spectrometry was performed on a Synapt XS instrument (Waters) operating in positive ionization mode with the following source settings: capillary voltage +3.0 and 2.0 kV, cone voltage 30 and 20 V, offset 4.0 V, desolvation temperature 600 °C, source temperature 150 °C, desolvation gas flow 800 L/h, cone gas flow 50 L/h, for MSe and MS/MS acquisition, respectively. MSe acquisition was performed in a mass range of 50-600 *m*/*z,* with collision energy ramp of 20-40 V. MS/MS acquisition was performed in targeted mode at *m/z* 160, with a collision energy ramp of 5–20 V and a scan time of 0.20 s. Mass calibration used sodium formate infused at 20 µL/min; lock-mass correction was applied using leucine-enkephalin (200 pg/µL in acetonitrile:water 50:50 v/v, 0.1% formic acid) acquired every 30 s via the lock-spray inlet. Data acquisition and processing were performed with MassLynx v4.2 (Waters).

Enzymatic reaction products and authentic standards were prepared at 20 µM in acetonitrile:water with 1 mM ammonium formate pH 9.70 (8:2, v/v). Samples were also spiked with 20 µM synthetic 2A3EA to confirm chromatographic co-elution. Metabolite identification was based on accurate mass within 5 ppm, isotopic pattern matching, MS/MS fragmentation profile, and retention time concordance with the synthetic standard of 2A3EA.

### Chromatographic analysis with Marfey’s reagent

A 100 μL aqueous solution of each analyte (0.3–10 mg/mL) was derivatized by the addition of NaHCO_3_ (40 μL, 1 M) and Marfey’s reagent (1-fluoro-2,4-dinitrophenyl-5-L-alanine amide) (200 μL, 1% w/v solution in acetone). Each mixture was incubated at 40 °C and 1000 rpm for 75 min. Upon cooling to room temperature, the reactions were quenched by the addition of HCl (20 μL, 2 N) and dried under a stream of nitrogen. The solid residue was resuspended in the HPLC eluent B (300 μL), centrifuged to eliminate some insoluble residue and submitted to HPLC analysis.

HPLC analyses were run by using an HPLC (VWR Hitachi Chromaster, Japan) equipped with a 5410 UV detector, a 5310 column oven, a 5260 auto sampler, and a 5110 pump. Separation was achieved on a Luna Omega Polar C18 column (250 x 4.6 mm, 5 μm) (Phenomenex, Bologna, Italy). The detection wavelength was set at 340 nm, with a flow rate of 1.2 mL/min and a column temperature of 35 °C. Injection volume was 5 μL. Two mobile phases were employed for sample elution: eluent A: CH_3_COONa 0.04 M pH 5.3/CH_3_CN (90:10 v/v) and eluent B: CH_3_COONa 0.04 M pH 5.3/CH_3_CN (50:50 v/v). The following elution program was performed: 0–20% B from 0 to 9 min (linear gradient); 20% B from 9.1 to 18 min (isocratic); 25-75% B from 18.1 to 24 min (linear gradient); 0% B at 24.1 (jump) and 0% B from 24.2 to 40 min (isocratic).

### Coevolutionary screening

Coevolutionary relationships among gene families were inferred using the cotr (correlated transitions) framework ^34^, which quantifies statistical associations based on correlated patterns of gene gain and loss across phylogenetically structured datasets. Coevolution was assessed using both cotr p-values and Jaccard similarity scores ^51^ computed from orthogroup presence/absence matrices.

Analyses were performed using orthogroup classifications from OrthoDB v12.2. Three datasets were considered: (i) a eukaryotic dataset filtered to include one representative per taxid and orthogroups present in at least 1% of genomes (57,684 orthogroups; 5,675 genomes; 69,829,543 genes); (ii) a metazoan dataset (41,047 orthogroups; 2,326 genomes; 36,457,013 genes); and (iii) a metazoan dataset excluding Aves (418 species). Orthogroups are defined by OrthoDB separately for each dataset; therefore, exclusion of specific taxonomic groups does not affect orthogroup construction but only the subset of genomes considered in the coevolution analysis.

Coevolution signals were used to identify candidate genes associated with tryptophan catabolism and to assess the evolutionary coherence of pathway components across eukaryotic lineages.

### Protein-ligand complex prediction

The AMR–NADH–2-AM ternary complex was predicted using a local installation of the open-source AlphaFold3 (V. 3.1) using a GPU node equipped with NVIDIA A100 80 GB. The AMR protein sequence (NP_001108070.1) and ligands SMILES (NADH and 2AM) retrieved from PubChem were used as input, with 2-AM protonation states assigned at pH 7.6 using Open Babel. The highest-ranked predicted complex was selected for downstream structural and mechanistic analysis. To assess reproducibility and sample conformational variability, 100 independent AlphaFold3 ^52^ runs were performed, each with a unique random seed (1-100, with 1 as reference). For each run, five structural models (model 0–4) were generated, and per-model confidence metrics (ipTM, pTM, PAE), were extracted from the AlphaFold3 JSON outputs. The five models within each seed run were ranked using the internal AlphaFold3 ranking score (Rank 1 corresponding to the highest score). 2-AM RMSD values were calculated for each predicted model relative to the reference model. All structural processing, metrics extraction, and plotting were performed using Python (v3.10) with pandas, numpy, and matplotlib, while molecular graphics were generated in PyMOL (v3.0).

### Analysis of the IPR006175 family in fungi and metazoa

Proteins belonging to the InterPro family IPR006175 (including bacterial AMD proteins) were retrieved from UniProtKB using the UniProt REST API by querying entries annotated with the InterPro cross-reference IPR006175 in a curated set of representative opisthokont species plus a limited set of bacterial species sequences included for comparison. Sequences were aligned to the Pfam hidden Markov model PF01042 using HMMER (hmmalign, --trim) to retain only columns corresponding to model match states, and the alignment was converted to A2M format using *sreformat*. Phylogenetic relationships were inferred using FastTree v2, with amino-acid distances computed using the BLOSUM45 substitution matrix. Tree inference was performed via a custom wrapper script that standardizes sequence identifiers before invoking FastTree The resulting phylogeny was visualized and annotated using Interactive Tree Of Life (iTOL).

### Phylogenetic analysis of ASPDH

The Maximum-Likelihood ASPDH tree was constructed with PhyML v3.0 (http://www.atgc-montpellier.fr/phyml/) (Guindon et al. 2010) using the automated model selection procedure (Lefort et al. 2017). Nodal support was estimated by bootstrap analysis with 100 replicates. Tree images were generated with FigTree v1.4.3 (https://github.com/rambaut/figtree/).

### Eukaryotic gene clusters identification and microsynteny analysis

A fungal gene cluster linking AMD to other genes of tryptophan degradation was first identified by querying EvolClustDB using the locus tag AFLA_079280 (UniProt: A0A7U2MQ13). To reconstruct the genomic loci with exon–intron structure, fungal homologues of ACMSD were retrieved by BLASTP searches against the NCBI RefSeq protein database (fungi) using the *Aspergillus clavatus* ACMSD (XP_001276020.1) as anchor (E-value ≤1×10⁻¹⁰). Non-redundant protein accessions were linked to genomic records via NCBI EDirect, and 20-kb windows (±10 kb) surrounding each coding sequence were retrieved in GenBank format. All CDS translations within each genomic window were extracted and queried locally against curated reference proteins (ACMSD, AMD, ALDH8A1 and HAAO) using BLASTP (E-value ≤1×10⁻⁵). Gene coordinates, exon structures and strand orientation were parsed from GenBank annotations to construct standardized synteny tables. When multiple loci were present in a species, the most gene-rich cluster was selected. Microsynteny was visualized using custom Python scripts, with gene classes color-coded and both exon-intron structure and transcriptional orientation explicitly represented.

### Comparative genomics of tryptophan catabolism genes

To improve orthogroup assignment accuracy, hidden Markov model (HMM)-based classification was used in place of high-throughput BLAST-based orthology inference, enabling more sensitive and specific detection of pathway components across divergent eukaryotic lineages. HMMsearch ^53^ (V. 3.4) was used to assign genes to orthogroups and to quantify their presence across genomes, generating normalized presence matrices ordered according to pathway structure. For gene families with complex paralogy (for example, ALDH8A1 and AMD/RIDA), competitive HMM models including out-of-family homologs were incorporated to improve discrimination between related enzymes. Taxonomic annotation was integrated using a curated classification of genomes.

Gene distributions across taxa were visualized at multiple taxonomic levels, and summary statistics were obtained by binarizing normalized scores using a fixed threshold (score > 0.3) and computing the fraction of genomes encoding each gene within predefined groups.

Phylogenetic patterns of gene retention and loss were further examined by mapping presence/absence data onto a representative species tree.

## Acknowledgment

We thank Tommaso Ganino, Barbara Prandi, Rebecca Ghidini, Martina Ferrarini, Simone Leggeri, Eva Percudani, Luca Calani, and Cristiano Negro for technical assistance, and Valérie de Crécy-Lagard for discussion. This work benefited from the equipment and framework of COMP-R initiatives, funded by the “Departments of Excellence” program of the Italian Ministry for University and Research (MUR, 2023-2027), from the European Research Council (ERC) under the European Union’s Horizon 2020 research and innovation program (PREDICT-CARE project, grant agreement 950050), and from the High Performance Computing facility of the University of Parma, Italy. This work was supported by the Telethon-Cariplo project GJC25F018 “Illuminating dark gene targets through coevolution” to RP and FS.

## References

1. Miura, H. et al. A link between stress and depression: Shifts in the balance between the kynurenine and serotonin pathways of tryptophan metabolism and the etiology and pathophysiology of depression. Stress 11, 198–209 (2008).

2. Le Floc’h, N., Otten, W. & Merlot, E. Tryptophan metabolism, from nutrition to potential therapeutic applications. Amino Acids 41, 1195–1205 (2011).

3. Schwarcz, R., Bruno, J. P., Muchowski, P. J. & Wu, H.-Q. Kynurenines in the mammalian brain: when physiology meets pathology. Nat. Rev. Neurosci. 13, 465–477 (2012).

4. O’Mahony, S. M., Clarke, G., Borre, Y. E., Dinan, T. G. & Cryan, J. F. Serotonin, tryptophan metabolism and the brain-gut-microbiome axis. Behav. Brain Res. 277, 32–48 (2015).

5. Cervenka, I., Agudelo, L. Z. & Ruas, J. L. Kynurenines: Tryptophan’s metabolites in exercise, inflammation, and mental health. Science 357, eaaf9794 (2017).

6. Xue, C. et al. Tryptophan metabolism in health and disease. Cell Metab. 35, 1304–1326 (2023).

7. Torrelli-Diljohn, A., Kulkarni, B. & Vitturi, D. A. Cell Signaling by Tryptophan Catabolism. Biochemistry 65, 1366–1394 (2026).

8. Canli, T. & Lesch, K.-P. Long story short: the serotonin transporter in emotion regulation and social cognition. Nat. Neurosci. 10, 1103–1109 (2007).

9. Mándi, Y. & Vécsei, L. The kynurenine system and immunoregulation. J. Neural Transm. 119, 197–209 (2012).

10. Giacomantonio, M. A. et al. Subversion of kynurenine-induced AHR activation in CD8 T cells by kynureninase-expressing antigen-presenting cells. Cell Rep. 45, 117149 (2026).

11. Okuda, S., Nishiyama, N., Saito, H. & Katsuki, H. Hydrogen peroxide-mediated neuronal cell death induced by an endogenous neurotoxin, 3-hydroxykynurenine. Proc. Natl. Acad. Sci. 93, 12553–12558 (1996).

12. Guillemin, G. J. Quinolinic acid, the inescapable neurotoxin. FEBS J. 279, 1356–1365 (2012).

13. Maitre, M., Taleb, O., Jeltsch-David, H., Klein, C. & Mensah-Nyagan, A. Xanthurenic acid: A role in brain intercellular signaling. J. Neurochem. 168, 2303–2315 (2024).

14. Duque, G. et al. Picolinic Acid, a Catabolite of Tryptophan, Has an Anabolic Effect on Bone In Vivo. J. Bone Miner. Res. 35, 2275–2288 (2020).

15. Morgan, T. H. Sex Limited Inheritance in *Drosophila*. Science 32, 120–122 (1910).

16. Figon, F. & Casas, J. Ommochromes in invertebrates: biochemistry and cell biology. Biol. Rev. 94, 156–183 (2019).

17. Van Hellemond, J. J., Klockiewicz, M., Gaasenbeek, C. P. H., Roos, M. H. & Tielens, A. G. M. Rhodoquinone and Complex II of the Electron Transport Chain in Anaerobically Functioning Eukaryotes. J. Biol. Chem. 270, 31065–31070 (1995).

18. Del Borrello, S. et al. Rhodoquinone biosynthesis in C. elegans requires precursors generated by the kynurenine pathway. eLife 8, e48165 (2019).

19. Nishizuka, Y. & Hayaishi, O. Enzymic Synthesis of Niacin Nucleotides from 1- Hydroxyanthranilic Acid in Mammalian Liver. J. Biol. Chem. 238, PC483–PC485 (1963).

20. Nishizuka, Y., Ichiyama, A., Gholson, R. K. & Hayaishi, O. Studies on the Metabolism of the Benzene Ring of Tryptophan in Mammalian Tissues. J. Biol. Chem. 240, 733–739 (1965).

21. Ichiyama, A. et al. Studies on the Metabolism of the Benzene Ring of Tryptophan in Mammalian Tissues. J. Biol. Chem. 240, 740–749 (1965).

22. Nishizuka, Y., Ichiyama, A. & Hayaishi, O. Metabolism of the benzene ring of tryptophan (mammals). in Methods in Enzymology vol. 17 463–491 (Elsevier, 1970).

23. Colabroy, K. L. & Begley, T. P. The Pyridine Ring of NAD Is Formed by a Nonenzymatic Pericyclic Reaction. J. Am. Chem. Soc. 127, 840–841 (2005).

24. Keys, L. D. & Hamilton, G. A. Mechanism for the conversion of alpha.-amino-.beta.-carboxymuconate.epsilon.-semialdehyde to quinolinate, an apparent nonenzmic step in the biosynthesis of the nicotinamide coenzymes from tryptophan. J. Am. Chem. Soc. 109, 2156–2163 (1987).

25. Li, T., Ma, J. K., Hosler, J. P., Davidson, V. L. & Liu, A. Detection of Transient Intermediates in the Metal-Dependent Nonoxidative Decarboxylation Catalyzed by α-Amino-β-Carboxymuconate-ε-Semialdehyde Decarboxylase. J. Am. Chem. Soc. 129, 9278–9279 (2007).

26. Davis, I., Yang, Y., Wherritt, D. & Liu, A. Reassignment of the human aldehyde dehydrogenase ALDH8A1 (ALDH12) to the kynurenine pathway in tryptophan catabolism. J. Biol. Chem. 293, 9594–9603 (2018).

27. Ragueneau, E. et al. The Reactome Knowledgebase 2026. Nucleic Acids Res. 54, D673–D681 (2026).

28. Kanehisa, M., Furumichi, M., Sato, Y., Matsuura, Y. & Ishiguro-Watanabe, M. KEGG: biological systems database as a model of the real world. Nucleic Acids Res. 53, D672–D677 (2025).

29. Caspi, R. et al. The MetaCyc database of metabolic pathways and enzymes and the BioCyc collection of Pathway/Genome Databases. Nucleic Acids Res. 42, D459–D471 (2014).

30. Colabroy, K. L. & Begley, T. P. Tryptophan Catabolism: Identification and Characterization of a New Degradative Pathway. J. Bacteriol. 187, 7866–7869 (2005).

31. Martins, T. M., Martins, C., Guedes, P. & Silva Pereira, C. Twists and Turns in the Salicylate Catabolism of *Aspergillus terreus*, Revealing New Roles of the 3-Hydroxyanthranilate Pathway. mSystems 6, e00230-20 (2021).

32. Malatesta, M., et al. C11orf54 catalyzes L-xylulose formation in human metabolism. Proc. Natl. Acad. Sci. 122, e2506597122 (2025).

33. Tsaban, T., et al. CladeOScope: functional interactions through the prism of clade-wise co-evolution. NAR Genomics Bioinforma. 3, lqab024 (2021).

34. Dembech, E., et al. Identification of hidden associations among eukaryotic genes through statistical analysis of coevolutionary transitions. Proc. Natl. Acad. Sci. 120, e2218329120 (2023).

35. Moi, D. & Dessimoz, C. Phylogenetic profiling in eukaryotes comes of age. Proc. Natl. Acad. Sci. U. S. A. 120, e2305013120 (2023).

36. Langschied, F. et al. Quest for Orthologs in the Era of Biodiversity Genomics. Genome Biol. Evol. 16, evae224 (2024).

37. Nishimura, Y., Omae, K., Tominaga, K. & Iwasaki, W. CORGIAS: identifying correlated gene pairs by considering evolutionary history in a large-scale prokaryotic genome dataset. NAR Genomics Bioinforma. 7, lqaf182 (2025).

38. Huttener, R. et al. Sequencing refractory regions in bird genomes are hotspots for accelerated protein evolution. BMC Ecol. Evol. 21, 176 (2021).

39. Yoneda, K., Sakuraba, H., Tsuge, H., Katunuma, N. & Ohshima, T. Crystal structure of archaeal highly thermostable L-aspartate dehydrogenase/NAD/citrate ternary complex. FEBS J. 274, 4315–4325 (2007).

40. Nasu, S., Wicks, F. D. & Gholson, R. K. The mammalian enzyme which replaces b protein of e. coli quinolinate synthetase is d-aspartate oxidase. Biochim. Biophys. Acta BBA - Protein Struct. Mol. Enzymol. 704, 240–252 (1982).

41. He, X., Kang, Y. & Chen, L. Identification of ASPDH as a novel NAADP-binding protein. Biochem. Biophys. Res. Commun. 621, 168–175 (2022).

42. Sethi, S., Martens, J. & Bhushan, R. Assessment and application of Marfey’s reagent and analogs in enantioseparation: a decade’s perspective. Biomed. Chromatogr. 35, e4990 (2021).

43. Allan, R. et al. Synthesis of Unsaturated Analogues of Glutamic Acid: Amination of Trianions From Unsaturated Dicarboxylic Acids With Chloramine. Aust. J. Chem. 49, 785–791 (1996).

44. Iwai, K. & Taguchi, H. DISTRIBUTION OF QUINOLINATE PHOSPHORIBOSYL-TRANSFERASE IN ANIMALS, PLANTS AND MICROORGANISMS. J. Nutr. Sci. Vitaminol. (Tokyo) 19, 491–499 (1973).

45. Ng, W.-K., Serrini, G., Zhang, Z. & Wilson, R. P. Niacin requirement and inability of tryptophan to act as a precursor of NAD+ in channel catfish, Ictalurus punctatus. Aquaculture 152, 273–285 (1997).

46. Fillgrove, K. L. & Anderson, V. E. The mechanism of dienoyl-CoA reduction by 2,4-dienoyl-CoA reductase is stepwise: observation of a dienolate intermediate. Biochemistry 40, 12412–12421 (2001).

47. Li, D., Li, M., Yan, M. & Xiong, Y. ASPDH inhibits the proliferation, migration, and invasion of liver cancer cells by regulating lactate metabolism and the NF-κB/PD-L1 pathway. Clinics 81, 100946 (2026).

48. Kumar, S. et al. TimeTree 5: An Expanded Resource for Species Divergence Times. Mol. Biol. Evol. 39, msac174 (2022).

49. He, Z. & Spain, J. C. Preparation of 2-aminomuconate from 2-aminophenol by coupled enzymatic dioxygenation and dehydrogenation reactions. J. Ind. Microbiol. Biotechnol. 23, 138–142 (1999).

50. Gassner, G., Wang, L., Batie, C. & Ballou, D. P. Reaction of Phthalate Dioxygenase Reductase with NADH and NAD: Kinetic and Spectral Characterization of Intermediates. Biochemistry 33, 12184–12193 (1994).

51. Glazko, G. V. & Mushegian, A. R. Detection of evolutionarily stable fragments of cellular pathways by hierarchical clustering of phyletic patterns. Genome Biol. 5, R32 (2004).

52. Abramson, J. et al. Accurate structure prediction of biomolecular interactions with AlphaFold 3. Nature 630, 493–500 (2024).

53. Eddy, S. R. Accelerated Profile HMM Searches. PLoS Comput. Biol. 7, e1002195 (2011).

