## Supplemental Figures S1-S9 for "Human ASPDH is a 2-aminomuconate reductase that produces L-2-aminohex-3-enedioic acid in tryptophan catabolism"

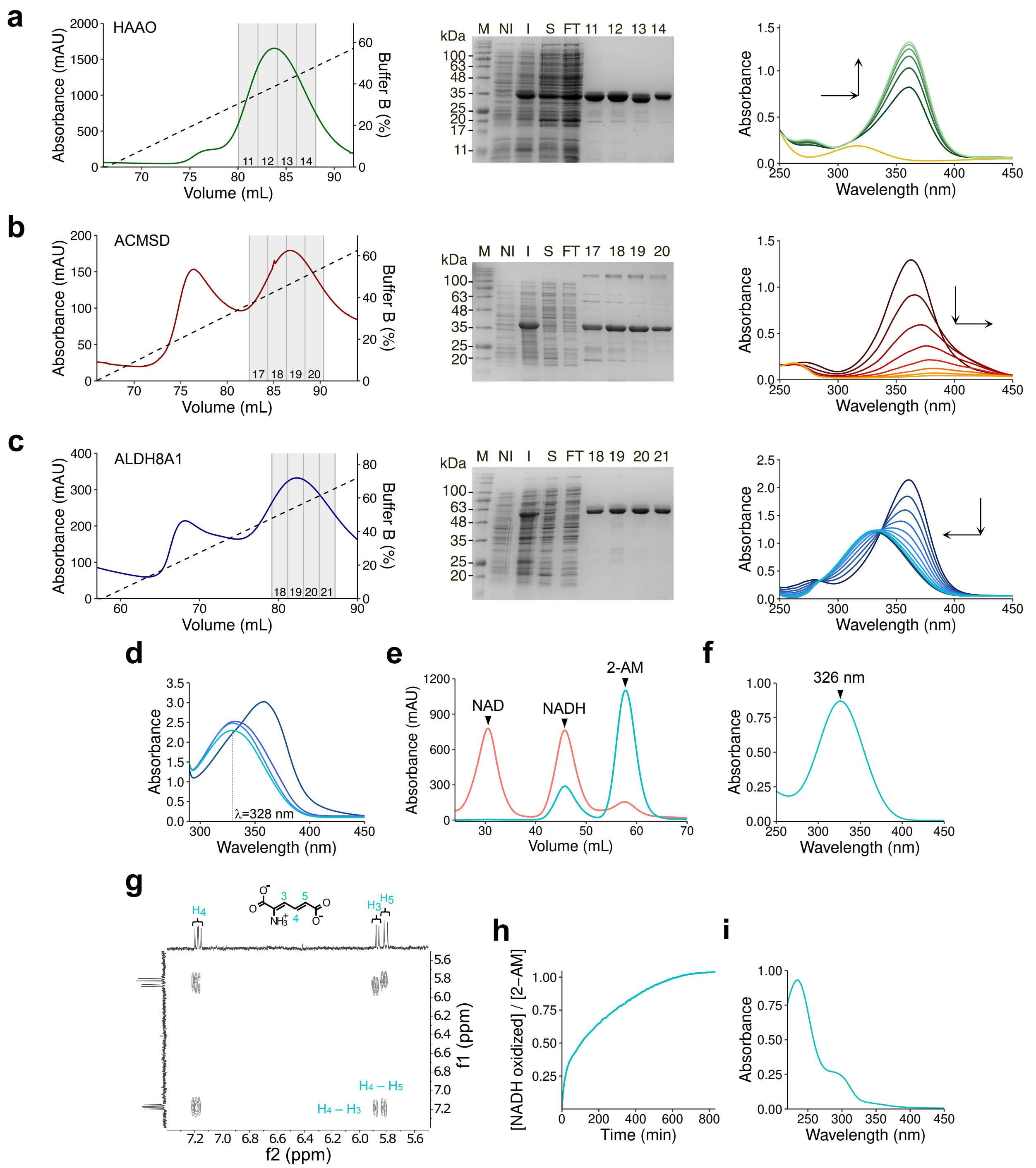


**Fig. S1 | Enzymatic synthesis and purification of 2AM.**

**a–c,** IMAC elution profiles (left) and SDS–PAGE gels (middle) showing purification of (a) HAAO, (b) ACMSD and (c) ALDH8A1. Lanes: marker (M), non-induced (NI), induced (I), supernatant (S), flow-through (FT), and numbered fractions, with matching fraction numbers between profiles and gels. UV–visible spectral changes (right) monitor substrate conversion by the purified enzymes. For HAAO, the first recorded spectrum (yellow) corresponds to 3-HAA prior to enzyme addition; subsequent spectra were acquired starting at 1 min after HAAO addition and every 30 s thereafter. For ACMSD and ALDH8A1 assays, upstream enzymes were included to generate substrates *in situ* (HAAO for ACMSD; HAAO and ACMSD for ALDH8A1), and reactions were initiated by addition of 3-HAA; spectra acquired every 30 s. **d**, UV–visible spectral changes during the one-pot enzymatic reaction leading to 2-AM formation. The reaction was initiated by addition of 3-HAA; spectra were acquired every 1 min, with the final spectrum recorded 2 min after the previous time point. **e**, Anion-exchange chromatography profile of the 2-AM formation reaction mixture. Absorbance at 260 (pinkish red) and 328 (cyan) nm is plotted versus elution volume. Peaks were assigned based on their UV–visible spectral properties. **f**, UV-visible absorption spectrum of purified 2-AM. **g**, COSY NMR spectrum of purified 2-AM, showing peaks at 7.15 (H4), 5.9 (H3), and 5.8 (H5) ppm. Proton assignments correspond to the chemical structure. **h**, NADH oxidation was monitored at 340 nm in the presence of glutamate dehydrogenase and α-ketoglutarate after addition of 2-AM, with time zero defined at 80 s. The ratio [NADH oxidized]/[2-AM] approaches unity consistent with a 1:1 stoichiometry between $NH_4^+$ release and NADH oxidation. **i**, UV–visible absorption spectrum of 2-AM after overnight incubation.

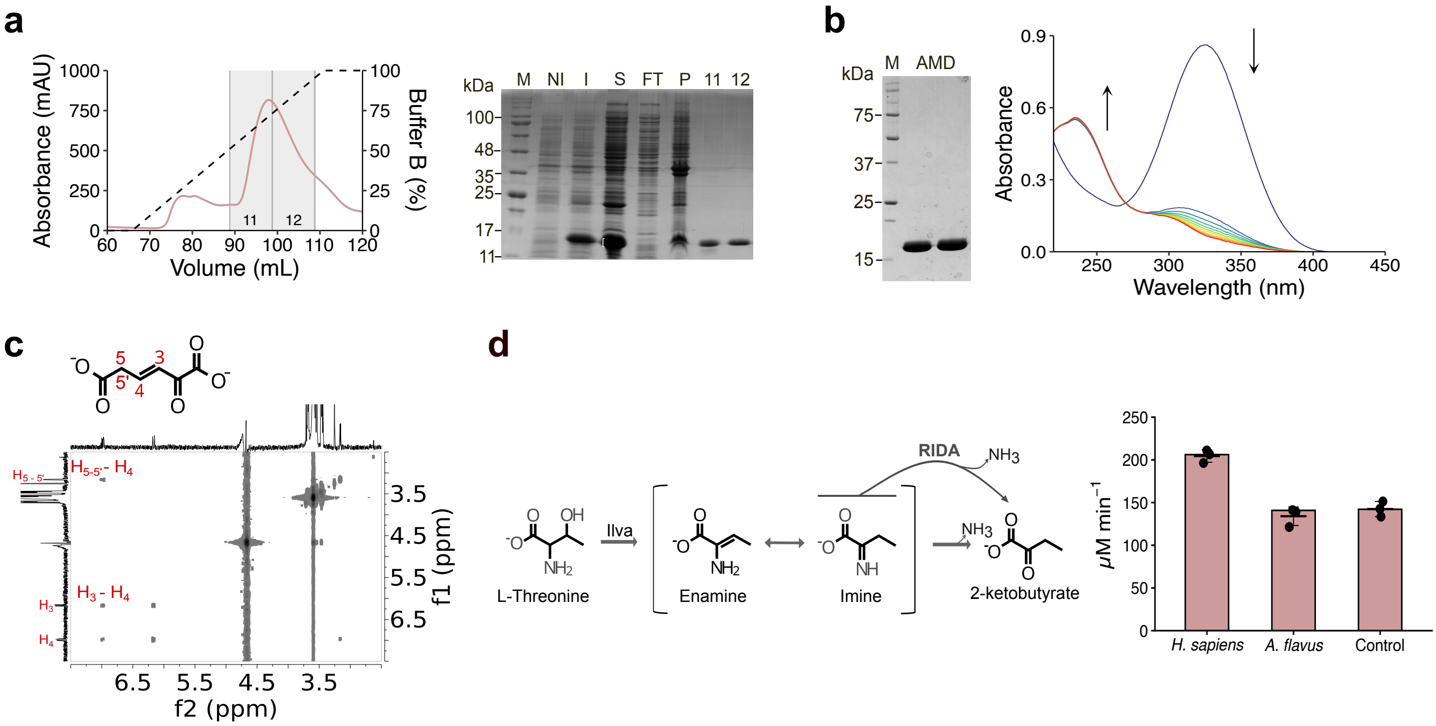

**Fig. S2 | Expression and functional characterization of AMD eukaryotic homologs.**

**a**, IMAC elution profile (left) and SDS–PAGE gel (right) showing purification of RIDA. Lanes: marker (M), non-induced (NI), induced (I), supernatant (S), flow-through (FT), pellet (P) and numbered fractions, with matching fraction numbers between profile and gel. **b**, SDS-PAGE gel (left) showing *A. flavus* AMD. Lanes: marker (M) and enzyme in duplicate (AMD). UV–visible spectral changes (right) following addition of *A. flavus* AMD to purified 2-AM in with an excess enzyme (0.5 μM), showing an enzyme-independent spectral transition. The first spectrum (dark blue) corresponds to 2-AM before enzyme addition; subsequent spectra were acquired starting at 5 s after AMD addition and every 30 s thereafter. **c**. COSY NMR spectrum of 2-oxohex-3-enedioate, obtained from purified 2-AM in presence of *A. flavus* AMD, showing peaks at 3.16 (H5-H5'), 6.95 (H4), and 6.20 (H3) ppm. Proton assignments are shown in the corresponding chemical structure. **d**, Schematic reaction (left) of RIDA family activity coupled with L-threonine dehydratase (Ilva) to produce 2-ketobutyrate. μM min-1 of NADH consumed (right) from RIDA activity in a three-enzyme coupled assay with GDH, Ilva and the enzymes from *A. flavus* (AMD) and *H. sapiens* (RIDA), and in a no-enzyme control. Points show individual measurements (n = 3); bars indicate the mean ± SD.

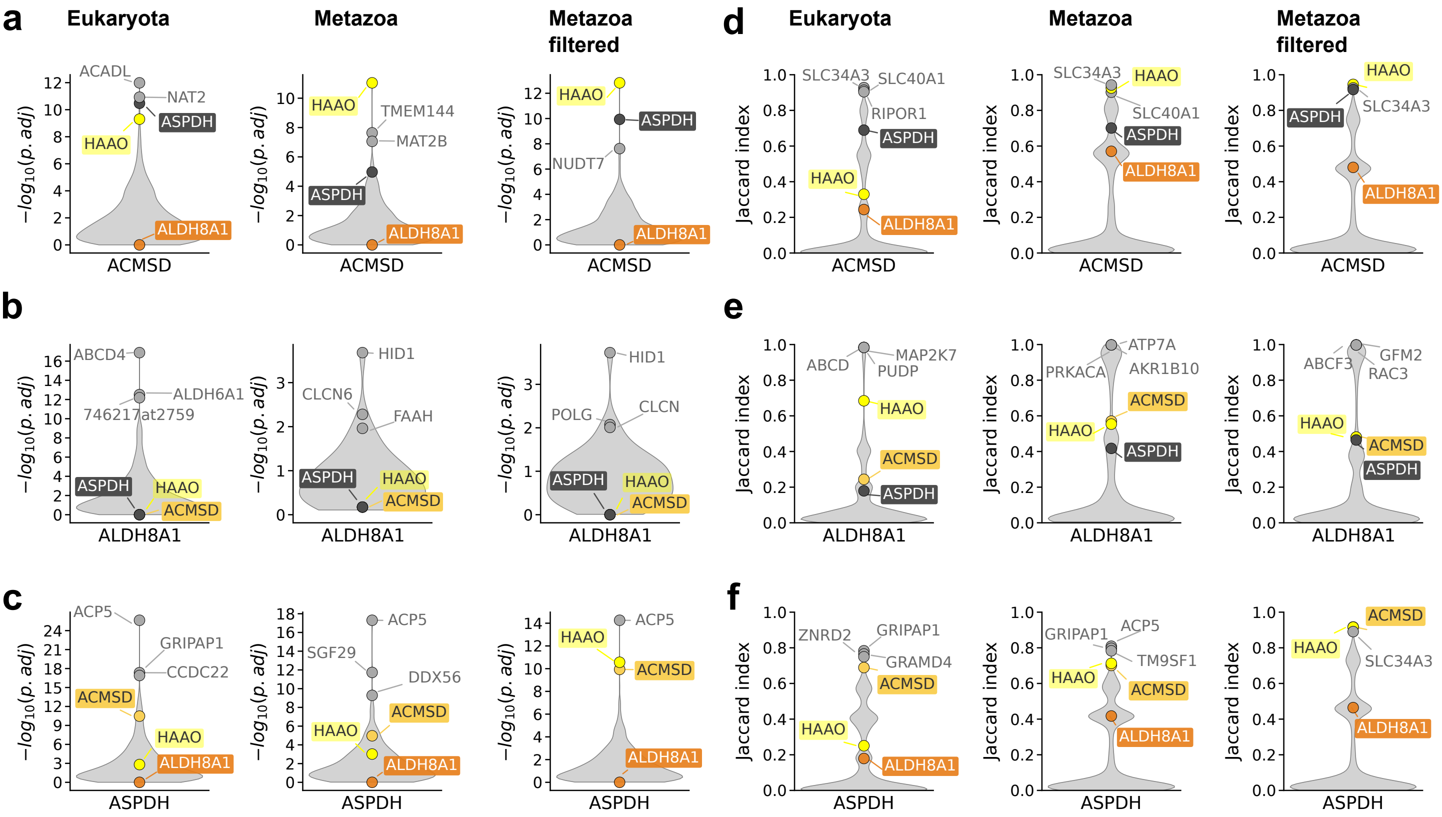


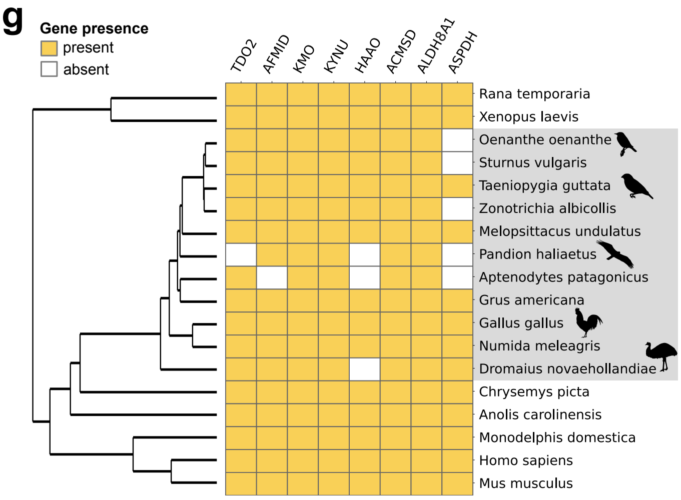


**h**

| Aves species | Accession number | % identity | E value | % GC |
|---|---|---|---|---|
| *G. gallus* | BG713440.1 | 97.89% | 2.00E-92 | 73 |
| *P. haliaetus* | GFNW01010924.1 | 75.09% | 4.00E-139 | 72 |
| *A. patagonicus* | HCAX01038676.1 | 76.07% | 5.00E-138 | 76 |
| *Z. albicollis* | GBBD01214159.1 | 60.00% | 3.00E-49 | 66 |
| *S. vulgaris* | GFDQ01005884.1 | 67.86% | 8.00E-122 | 70 |
| *O. oenanthe* | GFYT02009691.1 | 55.00% | 3.00E-77 | 72 |

**Fig S3 | Coevolutionary screening for KP genes.**

**a-f**, Violin plots showing the distributions of cotr P values (**a–c**) and Jaccard indices (**d–f**) obtained using ACMSD (**a**,**d**), ALDH8A1 (**b**,**e**), and ASPDH (**c**,**f**) as bait genes in eukaryotic (left; n species = 5,675, n genes = 77,042,201), metazoan (middle; n species = 2,326, n genes = 38,031,849), and metazoan excluding birds (right; n species = 1,908, n genes = 32,131,031) datasets. The three top genes, together with HAAO, ACMSD, ALDH8A1, and ASPDH, are indicated. Note that the inclusion of ALDH8A1 within a large orthogroup containing multiple aldehyde dehydrogenases obscures its coevolutionary relationships with other KP genes. **g**, Phylogenetic distribution of kynurenine pathway genes across selected Metazoa, with emphasis on Aves (grey shading). The heatmap indicates the presence (gold) or absence (white) of pathway enzymes across species ordered according to the rooted NCBI taxonomy tree. **h**, Table of selected avian species lacking annotated ASPDH protein sequences, for which corresponding transcript sequences were identified by tblastn using the *Gallus gallus* ASPDH protein (UniProt: A0A8V0X7L2_CHICK) as query. Columns report organism names, mRNA accession numbers (EST and TSA datasets), sequence identity (%), E-values relative to the *Gallus gallus* ASPDH protein, and mRNA G+C content.

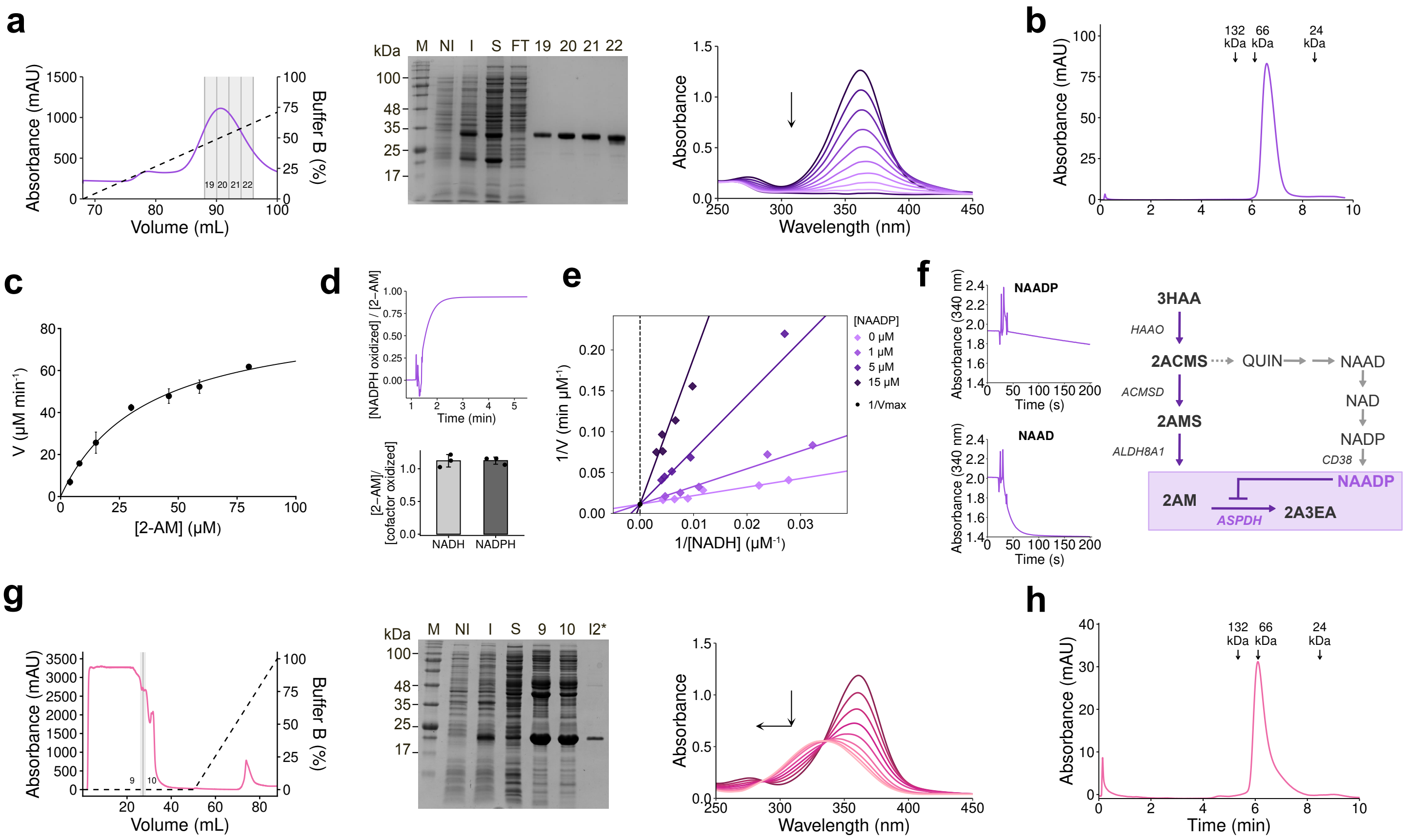

**Fig S4 | Human ASPDH functions as 2-aminomuconate reductase (2AMR).**

**a,** IMAC elution profiles (left), SDS–PAGE analysis (middle) and UV–visible spectral changes (right) for ASPDH canonical isoform (Uniprot A6ND91). Lanes: marker (M), non-induced (NI), induced (I), supernatant (S), flow-through (FT), and numbered fractions, with matching fraction numbers between chromatograms and gels. Upstream enzymes (HAAO, ACMSD and ALDH8A1) were included to generate substrates *in situ*, and reactions were initiated by addition of 3-HAA; spectra were acquired every 30 s. The decrease in the 2-AM signal is consistent with substrate turnover by ASPDH. **b,** SEC elution profile of ASPDH, with an estimated molecular mass of ~56 kDa. MW standards are indicated (BSA dimer, 132 kDa; BSA monomer, 66 kDa; trypsinogen, 24 kDa). **c,** Michaelis–Menten kinetics of ASPDH with 2-AM as substrate. Data points represent mean values (n = 3); error bars indicate ± SD; line indicates the best fit to the Michaelis–Menten equation. **d,** Top, NADPH oxidation at 340 nm was monitored in the presence of 2-AM following addition of ASPDH (~70 s). The [NADPH oxidized]/[2-AM] ratio approaches unity consistent with a 1:1 stoichiometry between NADPH oxidation and 2-AM reduction. Bottom, stoichiometry of 2-AM reduction relative to cofactor oxidation in ASPDH reactions containing NADH or NADPH. Bars indicate mean ± SD and dots indicate individual replicates (n = 3). **e,** Dependence of the initial rate of NADH oxidation and 2-AM reduction by ASPDH on increasing NAADP concentrations, showing inhibition of enzyme activity. Data points represent individual measurements and were fitted by global analysis using a competitive inhibition model. **f,** Time course of ASPDH activity (0.5 μM) at 340 nm using purified 2-AM in the presence of NADH (150 μM) and either NAAD (150 μM, top left) or NAADP (150 μM, bottom left). NAADP inhibited the reaction, whereas no inhibition was observed with NAAD. Right, schematic representation of the connection between tryptophan catabolism and NAD biosynthesis, highlighting NAADP, a NAD-derived metabolite, as an inhibitor of the ASPDH-mediated conversion of 2-AM to 2A3EA. **g,** IMAC elution profiles (left), SDS–PAGE analysis (middle) and UV–visible spectral changes (right) for ASPDH shorter isoform (Uniprot A6ND91-2), which eluted during the wash steps; the asterisk (*) indicates purified protein obtained after a second IMAC step. Under the same assay conditions as in (a), the 2-AM peak persisted, indicating lack of catalytic activity. **h,** SEC elution profile of ASPDH (shorter isoform), with an estimated molecular mass of ~69 kDa. MW standards are as in panel b.

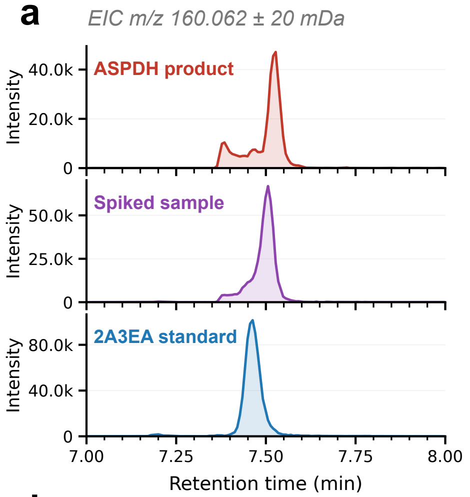


c

| m/z | Assignment | Δ ppm | Int. % |
|---|---|---|---|
| **160.0610** | **Parent ion $[M+H]^+$** | **-4.6** | **3.5** |
| 142.0483 | $-H_2O$ | -10.3 | 1.8 |
| 125.0253 | $-H_2O$, $-NH_3$ | +0.6 | 14.2 |
| 114.0555 | $-H_2O$, -CO | +0.5 | 100.0 |
| 101.0239 | $-H_2O$, -CO, $-CH_2$ | +0.6 | 34.8 |
| 100.0404 | $-C_2H_4O_2$ | +0.4 | 32.9 |
| 97.0301 | $-H_2O$, -CO, $-NH_3$ | +0.6 | 26.5 |
| 96.0418 | $-2(H_2O)$, -CO | +16.0 | 31.8 |
| 83.0135 | $-2(H_2O)$, -CO, $-NH_3$ | +1.0 | 38.6 |
| 82.0301 | $-C_2H_4O_2$, $-H_2O$ | +1.1 | 5.3 |
| 78.0343 | $-3(H_2O)$, -CO | +0.9 | 5.2 |
| 73.0286 | $-C_2H_4O_2$, -CO | +1.1 | 15.8 |
| 72.0434 | $-H_2O$, -CO, $COCH_2$ | +1.1 | 36.8 |
| 68.0502 | $-2(H_2O)$, -2(CO) | +0.3 | 10.2 |
| 55.0181 | $-H_2O$, -CO, $COCH_2$, $-NH_3$ | +1.6 | 3.2 |

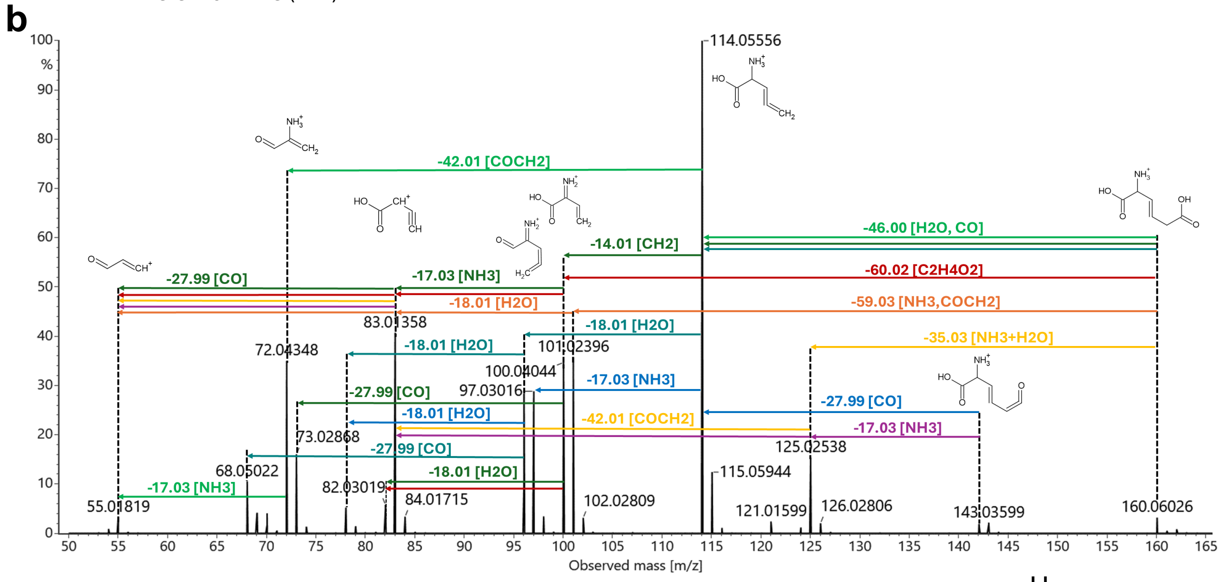


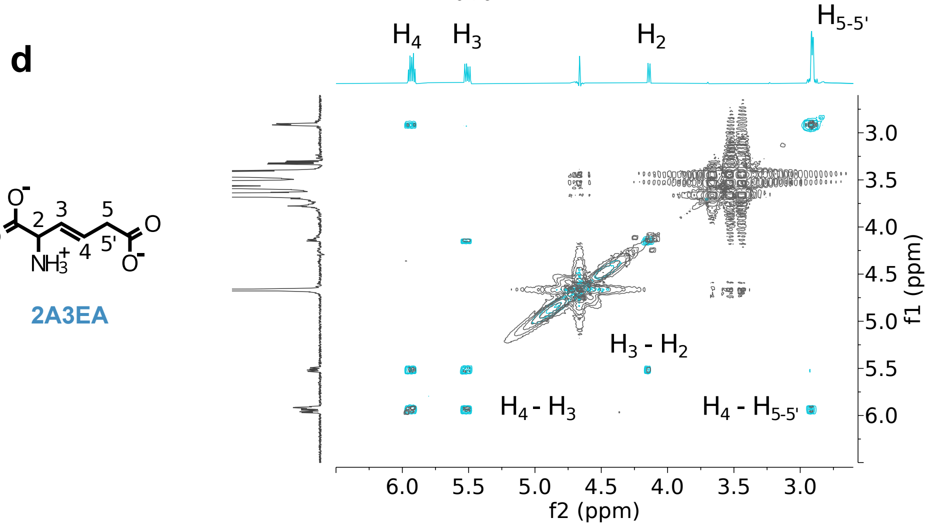


**Fig. S5 | Spectroscopic evidence for 2A3EA as the ASPDH reaction product.**

**a,** Detail of the extracted ion chromatogram (EIC) in the analyte elution window; expanded view of the retention time region 7.0–8.0 min showing the EIC at *m/z* 160.061 ± 20 mDa ($[M+H]^+$, $C_6H_{10}NO_4^+$) for three sample conditions. The spiked sample was prepared by addition of the reference standard to the sample matrix prior to analysis. **b,** MS/MS fragmentation map of 2A3EA**.** Positive-ion electrospray ionisation MS/MS spectrum was obtained using an optimised collision energy ramp. Relative intensities are normalised to the base peak (*m/z* 114.056, $[M+H-H_2O-CO]^+$, 100%). Coloured arrows indicate fragmentation pathways connecting precursor and product ions: dark green, −46.00 Da ($-H_2O$, −CO); red, −60.02 Da ($-C_2H_4O_2$); orange, −18.01 Da ($-H_2O$); teal, −17.03 Da ($-NH_3$); blue, −27.99 Da (−CO); light green, −14.01 Da ($-CH_2$); yellow, −35.03 Da ($-NH_3$ and $-H_2O$); purple, −59.03 Da ($-NH_3$ and $-COCH_2$); brown, −42.01 Da ($-COCH_2$). Proposed ionic structures are shown above the precursor and the major fragment ions. **c**, List of MS/MS fragments. Theoretical fragment *m/z* values, proposed neutral losses, mass errors relative to the closest centroid peak in the experimental spectrum (Δ ppm), and relative intensities are reported. **d,** Overlay of the COSY NMR spectra of the enzymatic product (black) and the 2A3EA standard (blue), with proton-coupling assignments.

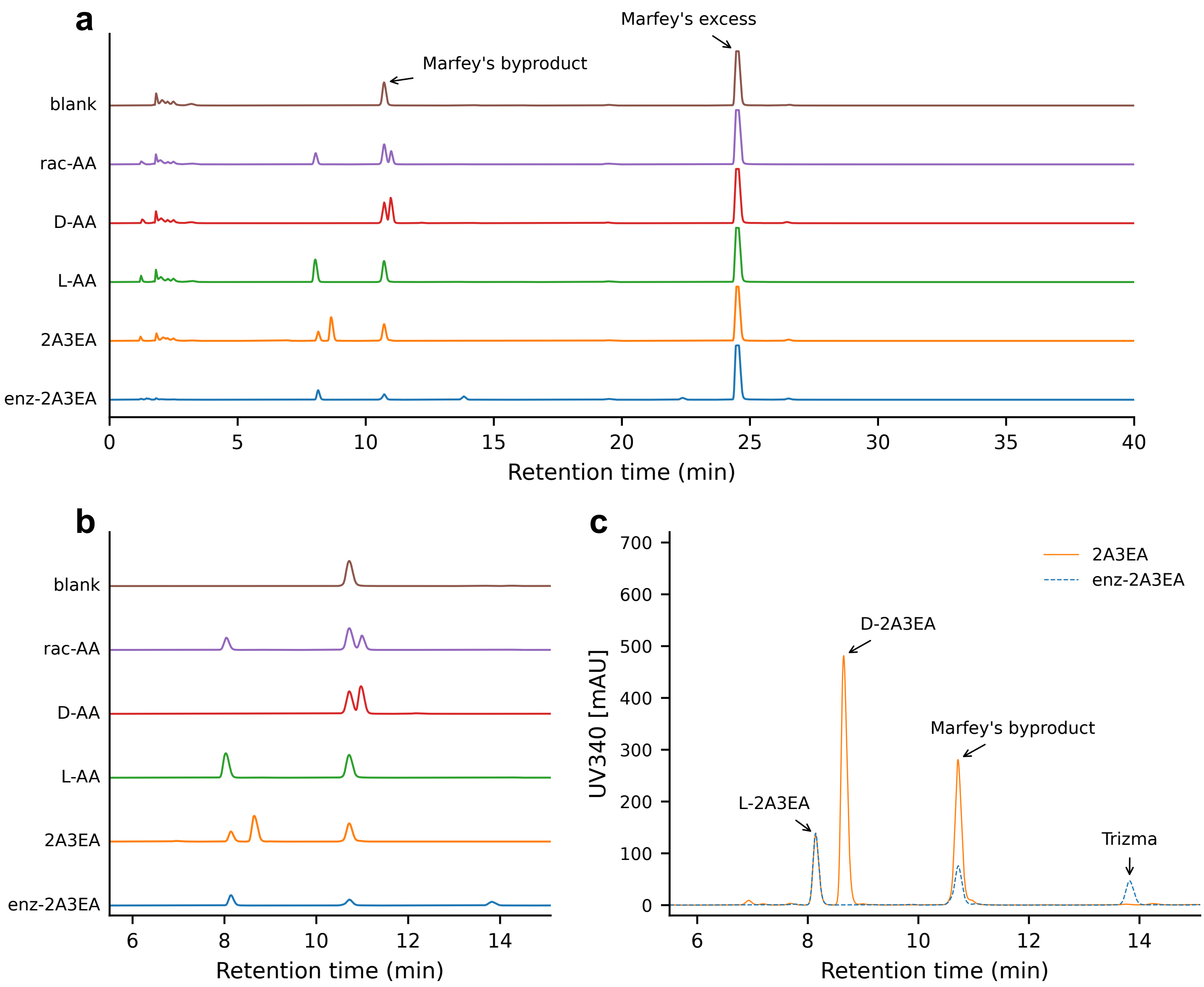


**Fig. S6 | Chromatographic analysis of 2A3EA enantiomers after derivatization with Marfey reagent.**

**a**, Stacked chromatograms of Marfey-derivatized standards and samples analysed by reversed-phase HPLC (340 nm). **b**, Expanded view (5.5–13.5 min). α-Aminoadipic acid (AA) (racemic, D and L) was used as a standard to validate chromatographic separation of diastereomers. The AA racemic standard resolves into two peaks that co-elute with the respective enantiopure standards. A chemically synthesized 2A3EA sample yields two peaks, whereas the enzymatic product (enz-2A3EA) yields a single peak. **c**, Overlay of 2A3EA and enz-2A3EA chromatograms on a common y scale highlighting the elution region of the derivatized product. Peaks assigned to L-2A3EA, D-2A3EA and Tris–HCl (Trizma) are indicated. Based on the established elution order of Marfey derivatives in reversed-phase chromatography, the enzymatic product is assigned the L configuration.

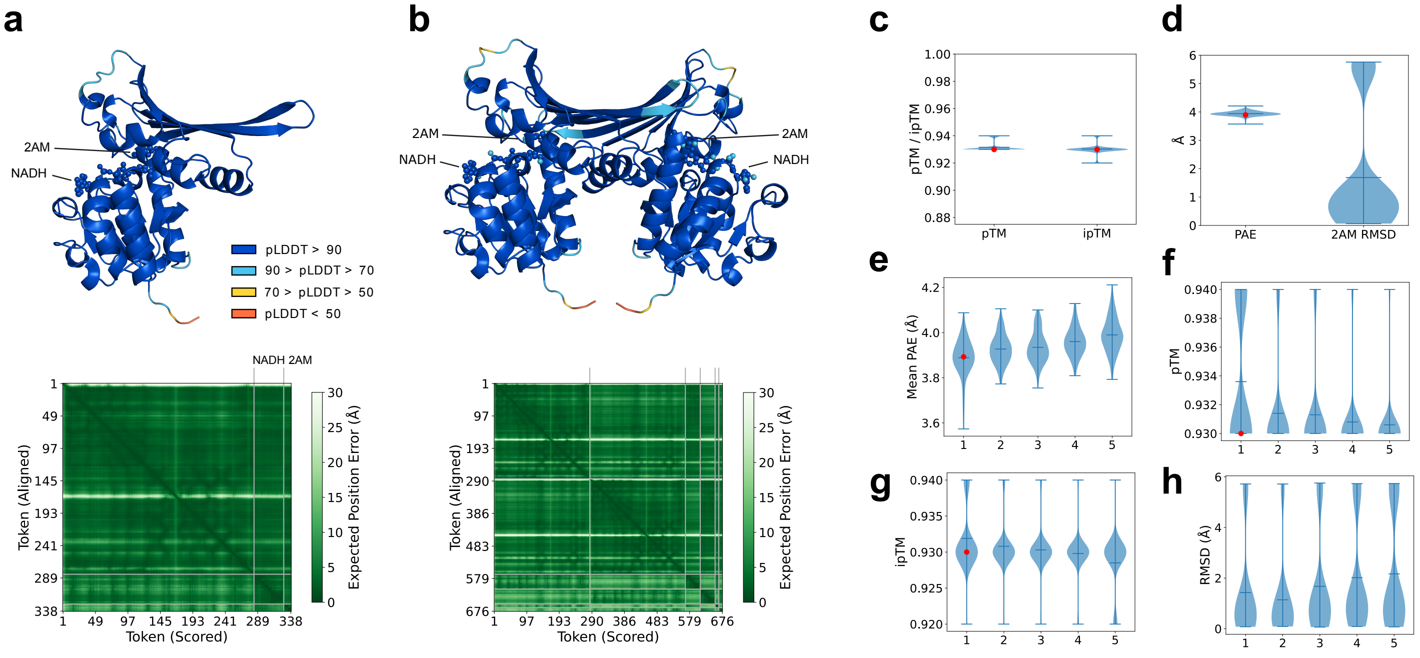


**Fig. S7 | AlphaFold3 protein-ligand complexes and confidence metrics.**

**a-b**, AlphaFold3 predictions of the monomeric and dimeric forms of AMR bound to NADH and 2AM, colored by per-residue (protein) or per-atom (ligands) pLDDT. Panels below display the corresponding predicted aligned error (PAE) plots of the reference model (seed1), indicating confidence in inter-residue distances. **c-d**, Violin plots of ipTM, pTM, mean PAE, and ligand RMSD across all models from 100 independent seed runs. Each violin shows the full distribution; central markers indicate the mean. The red dot indicates the reference model (seed1). **e-h**, Same metrics grouped by AlphaFold3 ranking score within each run (1 = highest ranking_score). The red dot indicates the reference model (seed1).

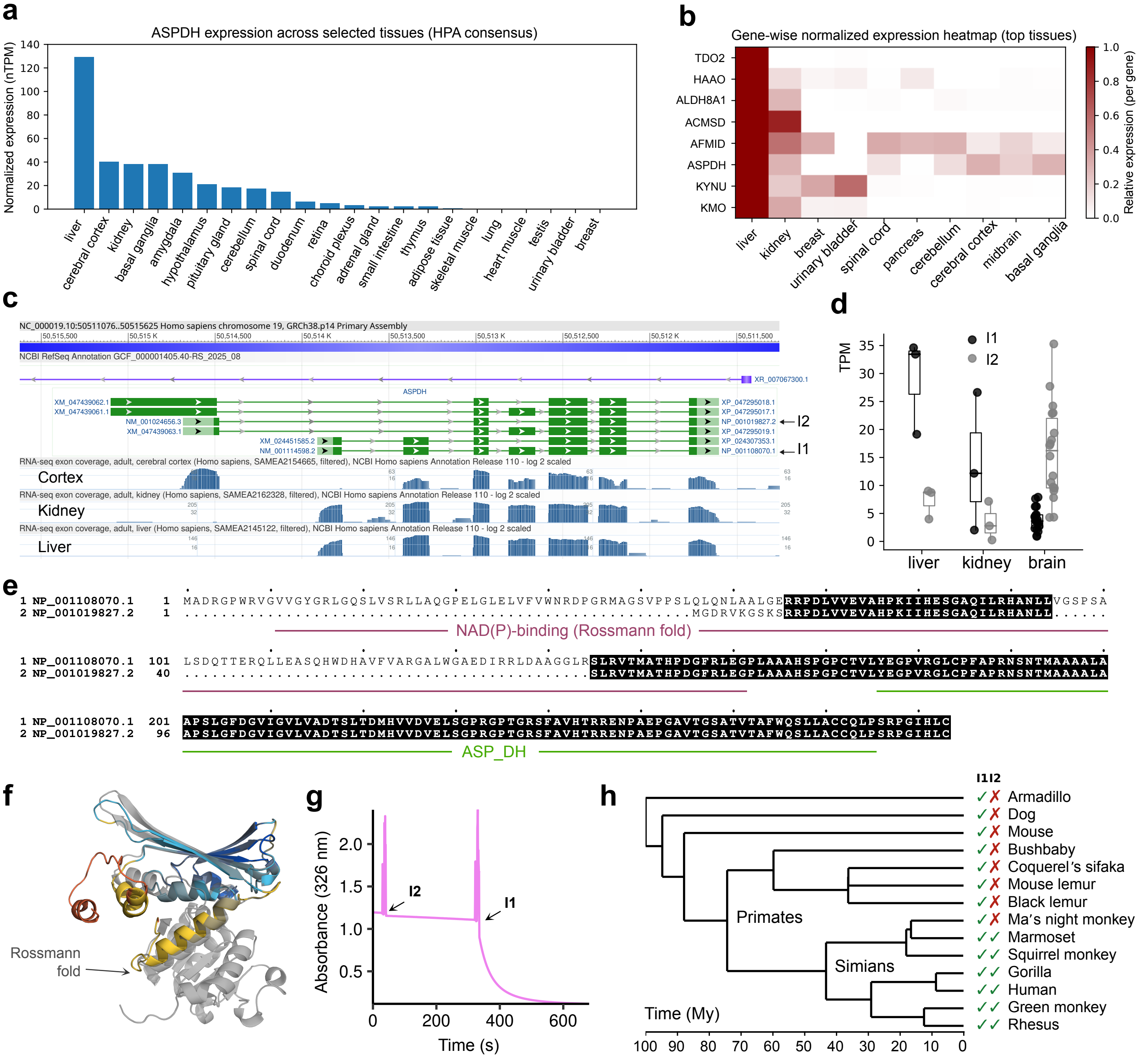


**Fig. S8 | Isoform expression and characterization of the human ASPDH locus.**

**a**, Tissue expression profile of ASPDH based on the Human Protein Atlas (HPA) consensus nTPM values. **b**, Gene-wise normalized heatmap of tryptophan catabolism genes across the ten tissues with highest average expression. **c**, NCBI view of the ASPDH locus showing curated isoforms 1 (I1) and 2 (I2) with RNA-seq tracks for cerebral cortex, kidney and liver. **d**, Kallisto-based quantification of I1 and I2 transcripts in brain, kidney and liver (ENA: PRJNA764684). **e**, Sequence alignment of I1 and I2 highlighting the boundaries of the NAD(P)-binding and aspartate dehydrogenase (ASP_DH) domains (InterPro). **f**, Superimposed AlphaFold3-predicted structures of ASPDH (grey) and the hypothetical I2 (coloured by per-residue pLDDT scores). **g**, Time course at 326 nm with purified 2-AM in the presence of NADH (60 μM) demonstrating lack of activity after addition of I2 (2 μM); addition of I1 (1 μM) as a control results in rapid signal decrease. **h**, TimeTree mammalian phylogeny showing species potentially encoding I1 and I2, inferred from comparative analysis of the genomic locus in the UCSC Genome Browser.

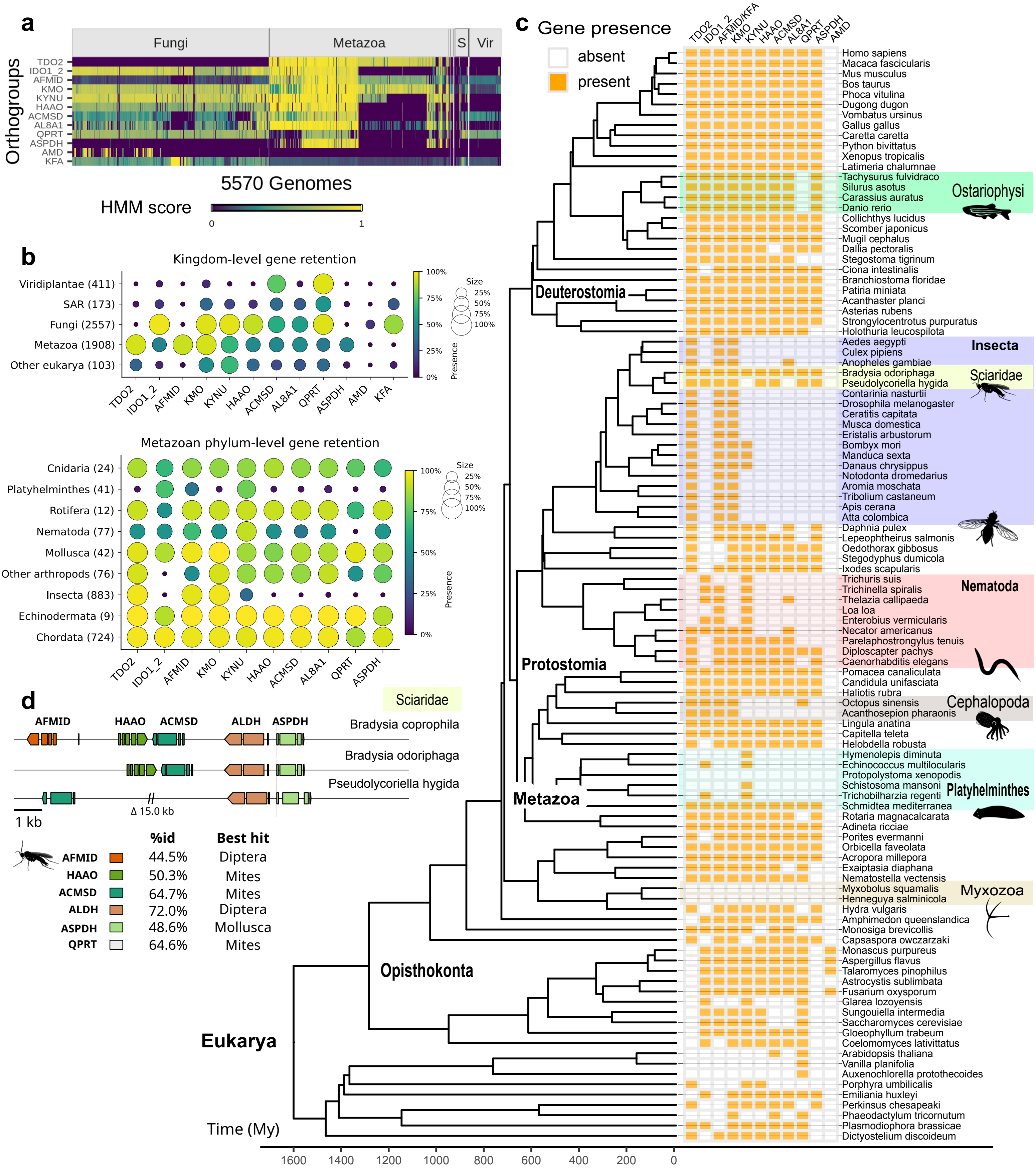


**Fig. S9 | Evolution of tryptophan catabolism via the kynurenine pathway in eukaryotes.**

**a,** Distribution heatmap of KP genes across 5,570 eukaryotic species. Gene presence (rows) was assessed across organisms (columns) using profile HMMs; colours indicate normalized hmmsearch scores. Taxonomic slices corresponding to Fungi, Metazoa, Sar (S), and Viridiplantae (Vir) are indicated. **b,** Fraction of species (Aves excluded) encoding each pathway enzyme across eukaryotic kingdoms (top) and metazoan phyla (bottom). Dot size and colour indicate the proportion of genomes with an assigned gene (threshold = 0.3; group sizes in parentheses). **c,** Phylogenetic distribution of pathway genes across selected eukaryotes with emphasis on Metazoa. The heatmap indicates presence (orange) or absence (white) of KP enzymes across a time-calibrated species tree inferred by maximum likelihood [34] and dated using TimeTree [48]. Major metazoan clades showing lineage-specific gene loss or retention are indicated. The tree is ladderized, with some nodes rotated to highlight independent loss and retention events. **d,** Microsynteny of the ASPDH locus in Sciaridae (Diptera), showing conserved genomic association with other KP genes. The table below reports the percentage identity and taxonomic origin of the best BLAST hit excluding Sciaridae. KFA, kynurenine-forming amidohydrolase; AL8A1 (ALDH8A1), aldehyde dehydrogenase 8 family member A1.
